# Imagined movement modulates cardiac-cortico-cortical and cardiac-cortico-cerebellar oscillatory networks

**DOI:** 10.1101/2025.10.21.683635

**Authors:** Diego Candia-Rivera, Mario Chavez, Fabrizio de Vico Fallani, Marie-Constance Corsi

## Abstract

Understanding the mechanisms of motor imagery, the mental simulation of movement without execution, is key for the development of neurotechnologies, including understanding inter-individual variability in motor imagery performance. For instance, for detecting covert motor intent in noncommunicative patients or refining motor commands through brain-computer interfaces. While motor imagery engages motor-related brain regions, its precise mechanisms remain unclear, particularly in relation to cardiac dynamics. Evidence suggests heart-rate variability features have potential to enhance tasks’ classifications, yet the brain-heart relationship is not well understood. In this study, we examined motor imagery learning using a task involving right-hand grasping imagery. We found that motor imagery is correlated with a task-dependent modulation of cardiac sympathetic activity and its relation with directed functional connectivity from the supplementary motor area to premotor and primary motor cortices. Additionally, cerebellar-supplementary motor area segregation, in relation to cardiac parasympathetic activity, indexed longitudinal motor learning. These results suggest that dynamic reconfiguration of brain-heart interactions contributes to sensorimotor function and learning-related physiology during motor imagery, supporting the brain-heart axis as a functional component of motor imagery rather than a passive correlate.

## Introduction

The physiological mechanisms underlying movement imagination without actual execution are of particular interest in many fields in relation with the neural computation of agency and movement (Ohata et al., 2020). Detecting the signatures of covert motor intent could enable important advances, ranging from the use of motor commands in brain-computer interfaces to clinical applications such as identifying awareness in non-communicative patients, supporting individuals with motor impairments, and enhancing rehabilitation strategies (Guerra et al., 2017; Khan et al., 2020; Owen et al., 2006; Zimmermann-Schlatter et al., 2008). In the context of brain–computer interfaces, understanding these mechanisms is also critical for explaining why motor imagery paradigms fail in a substantial proportion of users, a phenomenon commonly referred to as “illiteracy” or “non-responsiveness”. Research has shown that motor imagery engages distinct components of the motor cortex (Hanakawa et al., 2003; Henschke and Pakan, 2023; Kilintari et al., 2016). However, its underlying mechanisms remain unclear due to various confounding factors (Guger et al., 2003; Lebon et al., 2012), which likely contribute to the marked inter-individual variability observed in motor imagery performance, and robust non-invasive methods for capturing these mechanisms are still challenging (Edelman et al., 2024; Lebedev and Nicolelis, 2006; Lotte et al., 2018).

Motor imagery research largely focus on the primary motor cortex (M1) or its approximate location in scalp sensors (Berman et al., 2012; Dekleva et al., 2024; Facchini et al., 2002). This is because M1 coordinates with dorsal, ventral, and lateral premotor areas to orchestrate motor execution (Hoshi and Tanji, 2007), generating distinct activation patterns based on the limb involved (Lotze et al., 2000) and on the type of action executed (e.g., reaching vs grasping) (Cavina-Pratesi et al., 2010). M1 also acts as an integration hub within the somato-cognitive action network (Gordon et al., 2023). However, monitoring motor imagery solely through the primary motor cortex may not fully capture the underlying mechanisms. Since motor imagery involves learning the skill itself, the brain undergoes dynamic reorganization as these motor skills are acquired (Dayan and Cohen, 2011). Moreover, evidence points to the supplementary motor area (SMA) rather than M1 as leading the physiological dynamics underpinning motor imagery (Henschke and Pakan, 2023). A substantial body of neuroimaging research, including fMRI (Al-Wasity et al., 2021; Chen et al., 2009; Mehler et al., 2019) and PET studies(Malouin et al., 2003; Naito et al., 2002; Thobois et al., 2000), consistently demonstrates SMA changing during motor imagery, even in the absence of M1 activity. This is thought to reflect the SMA’s role in motor planning, sequencing, and internal generation of actions, functions that are central to imagining movements. In contrast, M1 is primarily responsible for the execution of voluntary movements, and its activation is typically subdued during motor imagery, as no actual movement is produced. Transcranial magnetic stimulation studies further support this distinction (Cona et al., 2017), showing that disruption of SMA impairs motor imagery performance, while M1 disruption does not significantly affect imagery. These findings suggest that the SMA serves as a key neural substrate for motor imagery by supporting the internal simulation of movement plans without triggering motor output.

Beyond central motor networks, motor imagery is also known to engage peripheral physiological processes, including autonomic changes. Interactions within the brain-heart axis are not static but are expected to be dynamic and context-dependent, with changes in task demands and learning altering the structure of brain-heart interactions. Importantly, such changes may be reflected in measures of statistical dependence without implying causal directionality. In line with this view, evidence suggests that incorporating heart-rate variability features into movement classifications have potential to enhance assessment accuracy (Fló et al., 2025; Georgaras and and Vourvopoulos, 2025; Kaufmann et al., 2012; Pfurtscheller et al., 2013; Scherer et al., 2007). However, the mechanisms linking cardiac dynamics with brain activity remain to be understood. These approaches, often considered within hybrid brain-computer interface systems, have so far failed to deliver robust solutions, likely due to their reliance on black-box machine learning implementations (Rajpura et al., 2024; Yang et al., 2022). From a broader perspective, motor imagery engages mechanisms underlying bodily self-consciousness (Candia-Rivera, 2026; Evans and Blanke, 2013), which are, in part, linked to brain-heart interactions (Candia-Rivera et al., 2024a; Park et al., 2016).

Given previous evidence highlighting the potential relevance of cardiac activity (Oishi et al., 2000; Peixoto Pinto et al., 2017) and brain oscillations primarily emerging from SMA (Ahn et al., 2013; Grosse-Wentrup and Schölkopf, 2012; Salardini et al., 2012; Shibasaki et al., 1993), we aimed at examining changes in the coupling between the cardiac sympathetic and parasympathetic indices (CSI and CPI) and directed connectivity in the alpha, beta and gamma bands from the SMA to three key motor-related regions: the cingulate motor cortex (CMC, comprising anterior and middle cingulate cortices (Dum and Strick, 1993)), the premotor cortex (PMC), M1 and cerebellum. We hypothesized that brain-heart mechanisms play a key role in motor imagery, by signaling changes in motor imagery accuracy and the functional plasticity associated with motor imagery learning (Figure 1a). Since visceral activity is strongly coupled with many motor-relevant regions at rest, including motor and cingulate cortices (Rebollo and Tallon-Baudry, 2022; Valenza et al., 2019), we expected motor imagery to be associated with a pronounced brain-heart segregation—similar to the functional disconnections observed within the cortical somato-motor hierarchy (Corsi et al., 2020).

**Figure 1.**
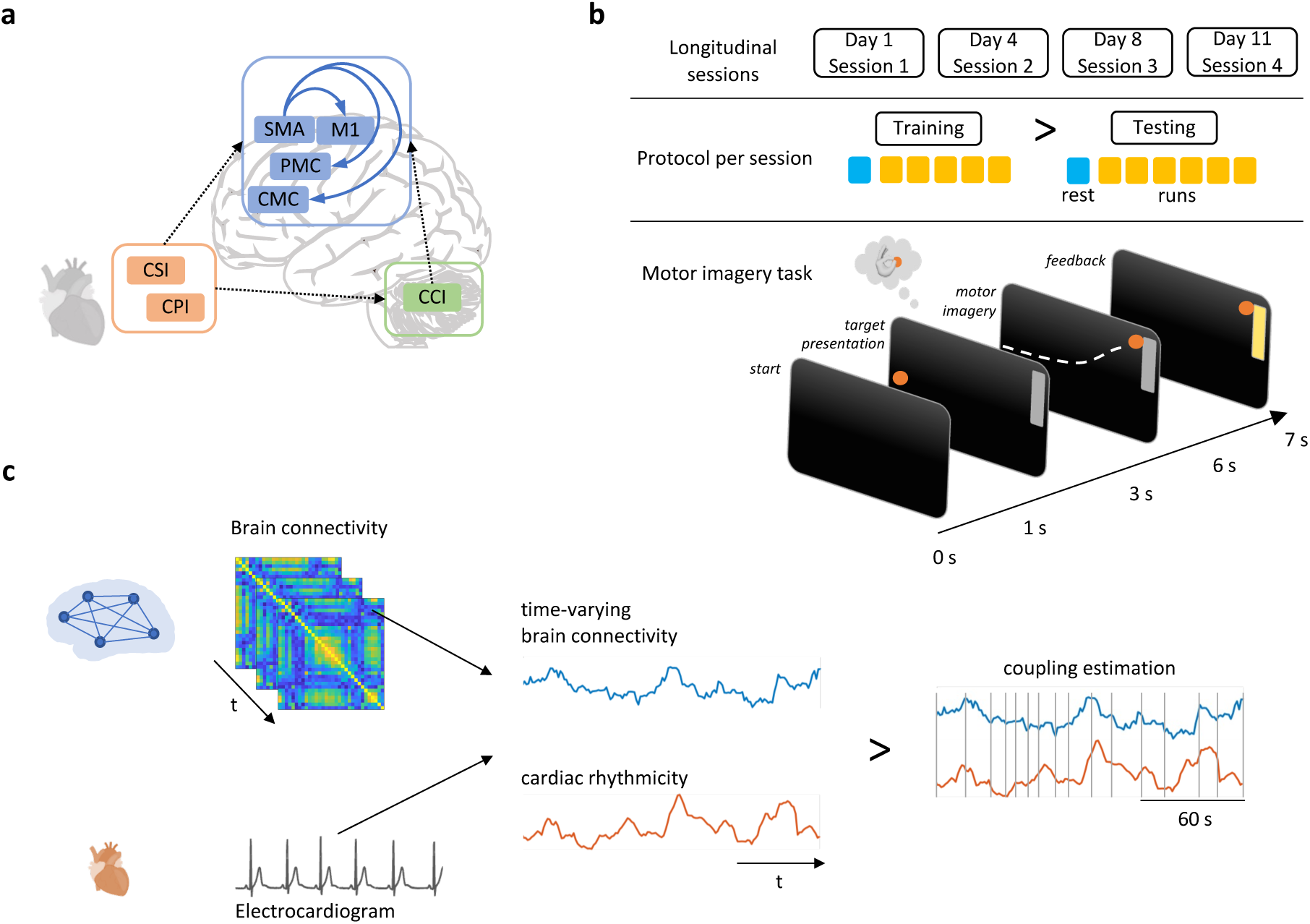
Experimental protocol and data processing pipeline. (a) The interaction between cardiac and brain components during motor imagery are studied. On the heart side, the cardiac sympathetic and parasympathetic indices (CSI and CPI). On the brain side, the supplementary motor area (SMA), primary motor cortex (M1), the premotor cortex (PMC), cingulate motor cortex (CMC), and the cerebellar crus I (CCI). (b) Participants completed four sessions of right-hand grasping motor imagery. Each session began with a resting-state period, followed by six runs of approximately three minutes each. In each run, participants performed 32 trials in a randomized order, imagining moving a target upward or by dropping the target by stopping their motor imagery. Participants were simultaneously recorded with MEG, EEG and ECG during the protocol (c) Data processing consisted of computing directed, time-varying brain connectivity in source-reconstructed MEG signals. Regions of interest included motor relevant brain areas, such as the Supplementary motor area, cingulate motor cortex, premotor cortex and the cerebellar crus.

In this study, we focus on a task consisting on subjects aiming to control the vertical position of a point on the screen (Wolpaw et al., 1991) (Figure 1b). To hit the target, the subjects performed a sustained motor imagery of right-hand grasping. Research employing this type of paradigm has unveiled alpha, beta and gamma activity, likely from SMA, is actively involved in motor imagery and motor planning (Ahn et al., 2013; Grosse-Wentrup and Schölkopf, 2012; Salardini et al., 2012; Shibasaki et al., 1993). Research on the cerebellum showed a leading role, especially in the motor learning process (Gao et al., 2018; Kipping et al., 2013; Stoodley et al., 2012). Reports on different markers of cardiac sympathetic activity showed a significant modulation during motor imagery (Oishi et al., 2000; Peixoto Pinto et al., 2017). While the exact specificity of these responses is still theorized, the link of those cardiac responses with those occurring in the brain remain uncertain.

We tested a recently proposed framework to study brain-heart interplay by quantifying the relationship between brain connectivity and estimators of cardiac sympathetic and parasympathetic activities (Candia-Rivera et al., 2024b) (Figure 1c). We found that motor imagery and its accuracy are correlated with cardiac sympathetic segregation with respect to the directed connectivity between SMA and other motor-related cortical regions. Additionally, we show that the change in the interaction between the cerebellum and SMA, in relation to cardiac parasympathetic activity, predicts motor imagery learning across sessions. Our findings highlight the potential role of cardiac rhythmicity in shaping connectivity patterns within motor-relevant cortical and cerebellar areas, underscoring the potential influence of interoceptive mechanisms in motor imagery.

## Methods

### Participants

This study comprises 20 subjects (aged 27.5 ± 4.0 years, 12 males and 8 females). Inclusion criteria were being right-handed and not presenting history of neurological or cardiovascular disease, and no previous experience in a motor imagery task. A written informed consent was obtained from subjects, according to the declaration of Helsinki. The local ethical committee approved this study (CPP-IDF-VI of Paris).

### Protocol

The task consisted of a two-target box task (Wolpaw et al., 2003) in which the subjects mentally aimed to control the vertical position of a cursor moving with constant velocity from the left to the right side of the screen. To hit the up-target, the subjects performed a sustained motor imagery of right-hand grasping and to hit the down-target they remained at rest-like condition in which they “dropped” the cursor. Each run consisted of 32 trials of approximately 5 s with up and down targets, consisting in a grey vertical bar displayed on the right part of the screen, equally and randomly distributed across trials.

The motor imagery accuracy values were gathered online (Corsi et al., 2020). In brief, in each session, a standard subject-specific predictive model was built in a training phase consisting in five consecutive runs without any feedback. The input features were based in the EEG power spectrum. The testing phase, used to gather the motor imagery accuracy consisted of six runs with visual feedback. The feedback consisted of a cursor that starts from the left-middle part of the screen and moves with a fixed velocity to the right part of the screen.

Although the sample size is modest (N = 20), the study employs a within-subject, repeated-measures design with multiple sessions and runs per participant, increasing sensitivity to condition- and session-related effects. Analyses focus on relative changes in brain-heart interaction patterns rather than absolute physiological amplitudes.

### Data acquisition

We carried out sessions with EEG signals transmitted to the BCI2000 toolbox(Schalk et al., 2004) via the Fieldtrip buffer (Oostenveld et al., 2011). Experiments were conducted with a 74 EEG-channel system (Easycap, Germany), referenced to the mastoid and grounded at the left scapula. MEG acquisition was performed with a system composed by 306 sensors (102 magnetometers and 204 gradiometers, Elekta Neuromag TRIUX MEG system). EEG and MEG signals were simultaneously recorded in a magnetic shielded room with a sampling frequency of 1 kHz and a bandwidth of 0.01–300 Hz. Individualized head position was acquired by Polhemus Fastrak digitizer system (Polhemus, Colchester, VT). Participants were seated in a comfortable and stable position with their arms placed on a support and their palms facing upwards.

Electromyogram signals were recorded from both arms. Electromyogram activity was visually inspected to ensure that participants were not moving during the recording sessions.

Individual head models were obtained after the fourth session using surface-based alignment(Gross et al., 2013) (T1 sequences, 256 sagittal slices, TR = 2.40 ms, TE = 2.22 ms, 0.80 mm isotropic voxels, 300x320 matrix; flip angle = 9°, 3T Siemens Magnetom PRISMA, Germany).

### Physiological data processing

### MEG data processing and source reconstruction

MRI images were preprocessed via the FreeSurfer toolbox (Fischl, 2012) and directly imported to the Brainstorm toolbox (Tadel et al., 2011). We used 15002 vertices projected to the automated anatomical labelling atlas for source reconstruction (Rolls et al., 2020). Source reconstruction was performed in line to previous pipeline aimed at gathering brain-heart relationships(Catrambone et al., 2024). Individual anatomy was used in each case. A three-shell sphere was created for each participant, and sources were computed using default parameters of the sLORETA method (Pascual-Marqui, 2002). The conductivities for surface estimation were set to default (scalp 1, skull 0.0125, brain 1). We employed a standard brain volume partition in which the brain volume is divided into a few regions of interest (ROI) defined according to anatomical and functional criteria, based on the current protocol. This allowed us to map 306 MEG sensors onto 18 ROIs from the AAL (see Table 4).

### Definition of regions and frequencies of interest

In this study investigating the brain-heart correlates of motor imagery and motor imagery learning, we focused on specific neural connections and frequency bands known to support motor-related functions and their modulation by autonomic signals. For motor imagery, we analyzed connectivity from the SMA to the PMC, M1, and CMC (Makoshi et al., 2011; Naito et al., 2002), specifically in the alpha, beta and low gamma bands (Ahn et al., 2013). These frequencies are associated with sensorimotor processing and fine motor control and are known to reflect top-down modulation and the maintenance of internal motor representations during imagery.

To investigate motor imagery learning, we examined connections from the cerebellar Crus I (Akkal et al., 2007), a region implicated in cognitive aspects of motor learning, to the SMA, M1, PMC and CMC (Solomon et al., 2021), focusing on the theta band, which has been linked to learning-related plasticity and coordination of long-range communication during task acquisition.

Additionally, we included the thalamus as a control region of interest, analyzing its connectivity focusing in the theta, alpha, beta, and gamma bands. The thalamus plays a central role in orchestrating motor imagery dynamics by gating sensorimotor signals and is also a key integrator of cardiac autonomic input, making it a critical hub for disentangling the interplay between neural and physiological components of motor imagery.

Source reconstruction was performed using individual anatomical MRIs for all participants, ensuring accurate forward modeling and head-sensor co-registration. A standardized inverse solution was employed, providing conservative and spatially smooth estimates that are commonly used for group-level MEG analyses. Given the known limitations of MEG sensitivity to deep brain structures, analyses were deliberately constrained to regions that can be reasonably captured with MEG. In the cerebellum, source reconstruction was restricted a priori to Crus I, which is among the most superficial and functionally well-characterized cerebellar regions accessible with MEG (Andersen et al., 2020; Stoodley and Schmahmann, 2009). Although the thalamus is a deep structure with limited MEG sensitivity, thalamic source estimates were included as secondary analyses and are reported in the Supplementary Material. These analyses were not used to support the main conclusions but were included to assess whether the brain-heart interaction effects observed in cortical and cerebellar regions could be retrieved in a deep structure with known localization limitations.

Although parietal and occipital regions have been frequently implicated in motor imagery (Decety, 1996), particularly in relation to visuospatial processing and mental simulation of movement, we deliberately excluded these areas to avoid confounding effects related to visual imagery and non-motor cognitive processes. Activation in these regions often reflects the visual or spatial components of imagined movements rather than motor preparation or execution per se, which could obscure the interpretation of motor-specific brain-heart interactions. Furthermore, occipital involvement may primarily indicate engagement of the visual cortex, which is highly sensitive to task instructions and individual imagery strategies, adding variability that is unrelated to the motor system or autonomic modulation. By focusing on the motor cortex—including SMA, PMC, M1, CMC, and cerebellum—we aimed to isolate the physiological and learning-related aspects of motor control, minimizing interference from higher-order cognitive or sensory processes. This targeted approach allows for a more specific investigation of motor-related dynamics and their specific relationship with cardiac activity.

### EEG data processing and motor imagery accuracy

Online features for motor imagery accuracy computation were based on EEG power spectra. Power spectra was estimated using an autoregressive model based on the Maximum Entropy Method (Kay and Marple, 1981) every 28 ms on time window of 0.5 s with a model order of 28. The classifier used was a Linear Discriminant Analysis. The classifier was independently trained per participant in each session, by searching the EEG channel and frequency bin with highest discrimination power. The training data was not used for further purposes.

Motor imagery accuracy values were obtained exclusively from the online BCI classifier, which was trained on EEG power spectrum features during separate training runs and tested on distinct runs. Importantly, MEG-derived brain connectivity, heart rate variability and brain-heart interaction measures were not used for classifier training or testing. All offline analyses relating brain-heart interaction to accuracy were therefore conducted post hoc and were fully independent of the online decoding procedure, preventing information leakage.

### Cardiac autonomic dynamics estimation

ECG time series were bandpass filtered using a fourth-order Butterworth filter, between 0.5 and 45 Hz. The R-peaks were identified in an automatized process based on the Pan-Tompkins algorithm (Pan and Tompkins, 1985). An automated detection of misdetections was implemented by searching sudden changes in the inter-beat intervals. The procedure was followed by a visual inspection of those potential misdetections. Finally, all the detected peaks were visually inspected over the original ECG, along with the inter-beat intervals histogram. Manual corrections of misdetections were performed if needed. Remarkably, manual corrections were minimal and occurred just in a few cases, indicative of the good signal to noise ratio of the dataset.

The cardiac sympathetic and parasympathetic activities were estimated through a method based on the time-varying geometry of the inter-beat interval Poincaré plot(Candia-Rivera et al., 2025c). The method combines ongoing changes in heart rate and heart rate variability to provide the Cardiac Parasympathetic Index (CPI) and Cardiac Sympathetic Index (CSI). It is worth mentioning that these markers were appropriately validated in standard autonomic elicitation, demonstrating their accuracy and robustness for a time-resolved depiction of cardiac autonomic dynamics.

For a detailed explanation of the cardiac autonomic estimation, refer to the original method article (Candia-Rivera et al., 2025c).

### Brain-heart coupling estimation

Source reconstructed MEG spectrogram was computed using the short-time Fourier transform with a Hanning taper. Calculations were performed through a sliding time window of 2 seconds with a 50% overlap, resulting in a spectrogram resolution of 1 second and 0.5 Hz. Time series were integrated within three frequency bands (theta: 4-7.5 Hz, alpha: 8-11.5 Hz, beta: 12-29.5 Hz, gamma: 30-45 Hz).

The directed time-varying connectivity between power series of two regions was quantified using autoregressive model gathering linear relationships with a 15 s sliding time window (Candia-Rivera et al., 2024b), which was aimed for capturing the directed temporal dynamics of a brain region, by leveraging on the dependency of its past values and another brain region, represented as the external term in the model. The connectivity model was set to minimize the adjusted error using least squares. The directed, time-varying connectivity is obtained from the adjusted coefficient from the external term.

Information-theoretic measures were used to quantify statistical dependencies between neural and cardiac signals. These measures characterize associations and changes in interaction structure but do not provide information about causal direction or statistical influence. Brain-heart coupling was quantified by considering the relationships between brain connectivity fluctuations and CPI or CSI. The brain-heart coupling was assessed using Maximal Information Coefficient (MIC) (Reshef et al., 2011), a method that quantifies the coupling between two time series at an adapted time scale that maximizes the mutual information, normalized by the minimum joint entropy, resulting in an index in the range 0-1.

MIC was computed using the standard implementation proposed by Reshef et al. (2011). Alpha exponent was set to 0.6 and parameter *c* (determining how many more clumps there will be than columns in every partition was set to 10), without manual tuning of grid resolution or smoothing parameters. Importantly, MIC was used as a descriptive measure of statistical dependence and not as a predictive or fitting model. Brain-heart interaction values were computed independently for each run and session, and no information from accuracy measures or linear model outcomes was used in the estimation of MIC, thereby avoiding circularity.

For a detailed explanation of the brain-heart coupling estimation, refer to the original method article (Candia-Rivera et al., 2024b).

### Control for physiological confounds

To assess potential contamination of MEG signals by muscle activity, we quantified frequency-domain coupling between MEG sources and simultaneously recorded EMG signals from the left and right arms. Spectral estimates were computed using a multi-taper fast Fourier transform approach. For each trial, Fourier spectra were estimated using a frequency smoothing of ±2 Hz, retaining single-trial estimates to enable coherence computation. Analyses focused on the high-frequency range (30-100 Hz; omitting the range 49-51 Hz of line noise), where EMG-related contamination is most prominent. Magnitude-squared coherence was then computed between MEG sources of interest and the two EMG channels. The resulting coherence spectra were inspected for broadband, zero-lag coupling patterns characteristic of EMG leakage.

To evaluate whether cardiac indices were primarily driven by respiratory dynamics, we conducted additional sensitivity analyses relating cardiac indices (CSI and CPI) to respiration. Breathing phase patterns were extracted from the ECG envelope (O’Brien and Heneghan, 2007), then breathing rates were computed using a simple peak detection algorithm on the phase signals. Spearman correlation analyses were performed between these respiratory measures and the cardiac indices across trials. This analysis allowed us to test whether variations in CSI and CPI could be explained by changes in breathing rate, rather than reflecting independent cardiac autonomic dynamics, as previously reported in resting state (Candia-Rivera and Chavez, 2025).

### Statistical analysis

Statistical analyses were primarily based on GLMs to identify the brain regions that best predicted the modeled variables. Region selection was performed by iteratively fitting separate models, each time varying the ROIs, and selecting the region with the lowest p-value for the effect of interest. This region-selection procedure was performed exclusively on brain-heart interaction metrics and was independent of the online motor imagery classifier, ensuring that feature selection did not exploit performance-related information.

The first GLM aimed to identify brain-heart dynamics consistently modulated by motor imagery. The model was defined as: *Δ Brain-Heart ∼ 1 + (1 | Participant).* This tested whether changes in brain-heart coupling (*ΔBrain-Heart)* from rest were significantly different from zero, as indicated by the p-value of the intercept.

The second GLM assessed brain-heart dynamics associated with variations in motor imagery accuracy. The model was: *Accuracy ∼ Δ Brain-Heart + (1 | Participant)*. Here, the p-value corresponding to the Δ Brain-Heart term was reported.

The third GLM focused on longitudinal motor imagery learning effects. The model was specified as: *Accuracy ∼ Session + ΔBrain-Heart + ΔBrain-Heart × Session + (1 | Participant)*. In this case, the interaction term Δ Brain-Heart*×*Session was evaluated, and its p-value reported. All GLMs included random intercepts for participants to account for inter-individual baseline differences, but also inter-individual differences in their learning rates (Sitaram et al., 2017). Random slopes were not included due to the modest sample size and to avoid model overparameterization. No additional covariates (e.g., age or sex) were included, as the study was not powered for subgroup analyses.

P-values from the GLMs were corrected for multiple comparisons using the Benjamini–Hochberg procedure to control the false discovery rate (FDR).

Post-hoc analyses were conducted on the brain regions identified by the GLMs, using non-parametric statistics. First, we tested whether the brain connection identified in the first GLM consistently changed its coupling with CSI or CPI across all sessions and motor imagery runs. For this, we compared the initial resting-state period of each session against each run using a two-tailed non-parametric Mann-Whitney test for paired comparisons, with significance set at p < 0.05.

Second, we examined whether changes in the coupling between the brain connection identified in the second GLM and CSI or CPI consistently correlated with motor imagery accuracy scores. This was assessed using Spearman correlation, with significance also set at p < 0.05. In addition, we performed a repeated measures version of the Spearman correlation. This approach accounts for the non-independence of repeated observations within participants by first applying rank transformations within each subject and then removing subject-specific means to obtain residuals. The correlation coefficient was calculated on these residuals, providing an estimate of the within-subject Spearman correlation. To assess statistical significance, a permutation procedure (10,000 iterations) was performed, in which residuals were randomly permuted within subjects to generate a null distribution of correlation values. Two-tailed permutation-based p-values were then derived by comparing the observed correlation with this null distribution.

Third, we tested whether the change in coupling between the brain link identified in the third GLM and either CSI or CPI increasingly correlated with motor imagery accuracy across sessions, as a potential signature of motor imagery learning. For each session, we averaged the accuracy across runs and computed its Spearman correlation with the corresponding average change in brain-heart coupling. This analysis was performed across all four sessions and confirmed that the correlation coefficients increased over time.

All post-hoc analyses were also replicated using brain connectivity and CSI or CPI alone, and the results were reported alongside those from the brain-heart coupling analyses. Additional analyses conducted on the opposite hemispheres and the thalamus are reported in the supplementary material.

## Results

Twenty healthy participants, without prior experience in motor imagery tasks, completed four sessions of motor imagery training. Each session included a resting-state recording followed by six runs of a motor imagery task, making a total of 480 individual motor imagery recordings. During the task, participants aimed to control the vertical position of a cursor on the screen by performing sustained motor imagery of right-hand grasping (Figure 1b). Physiological recordings included an initial anatomical MRI for subsequent source reconstruction, as well as EEG, ECG, and MEG during the experiment. EEG was used to train an individualized motor imagery classifier, providing an online accuracy and feedback, while MEG and ECG allowed for an a posteriori analysis of the physiological dynamics underlying motor imagery.

### Brain-heart dynamics are modulated by motor imagery

Our analysis focused on the coupling between cardiac rhythmicity and connectivity patterns in motor-related areas, derived from source-reconstructed MEG signals. First, we identified brain-heart coupling patterns that characterized the motor imagery condition and those that tracked motor imagery accuracy across the 480 recordings. We examined the coupling between the cardiac indices (CSI and CPI) and directed connectivity in the alpha, beta and gamma bands from the SMA to three key motor-related regions (CMC, PMC and M1). The computation of brain-heart dynamics was based on our recent approach to characterize cardiac-cortico-cortical dynamics(Candia-Rivera et al., 2024b). In brief, it incorporates the time-resolved estimation of heart rate and rhythms to derive CSI and CPI(Candia-Rivera et al., 2025c) and linear relationship between power series, modeled in an autoregressive process, to derive directed connectivity measures. Then, the coupling between CSI or CPI and directed connectivity series was derived from information-theoretic estimations to capture potential non-linear patterns (see Methods for detail).

We used generalized linear models (GLMs) to identify triads involving the CSI/CPI, SMA, and other motor areas. Our goal was to determine the physiological dynamics that are consistently modulated across most motor imagery sessions and runs. Specifically, the model assessed differences in brain-heart interactions between resting state and motor imagery, identifying changes that were significantly different from zero. As shown in Table 1, motor imagery significantly modulated the coupling between the CSI and the directed connectivity from the SMA to all three motor areas, in alpha, beta and gamma bands, in contralateral and ipsilateral dynamics. Noteworthy, the one that presented a stronger modulation is the coupling between CSI and ipsilateral brain connectivity from SMA to CMC in the beta band.

**Table 1.**
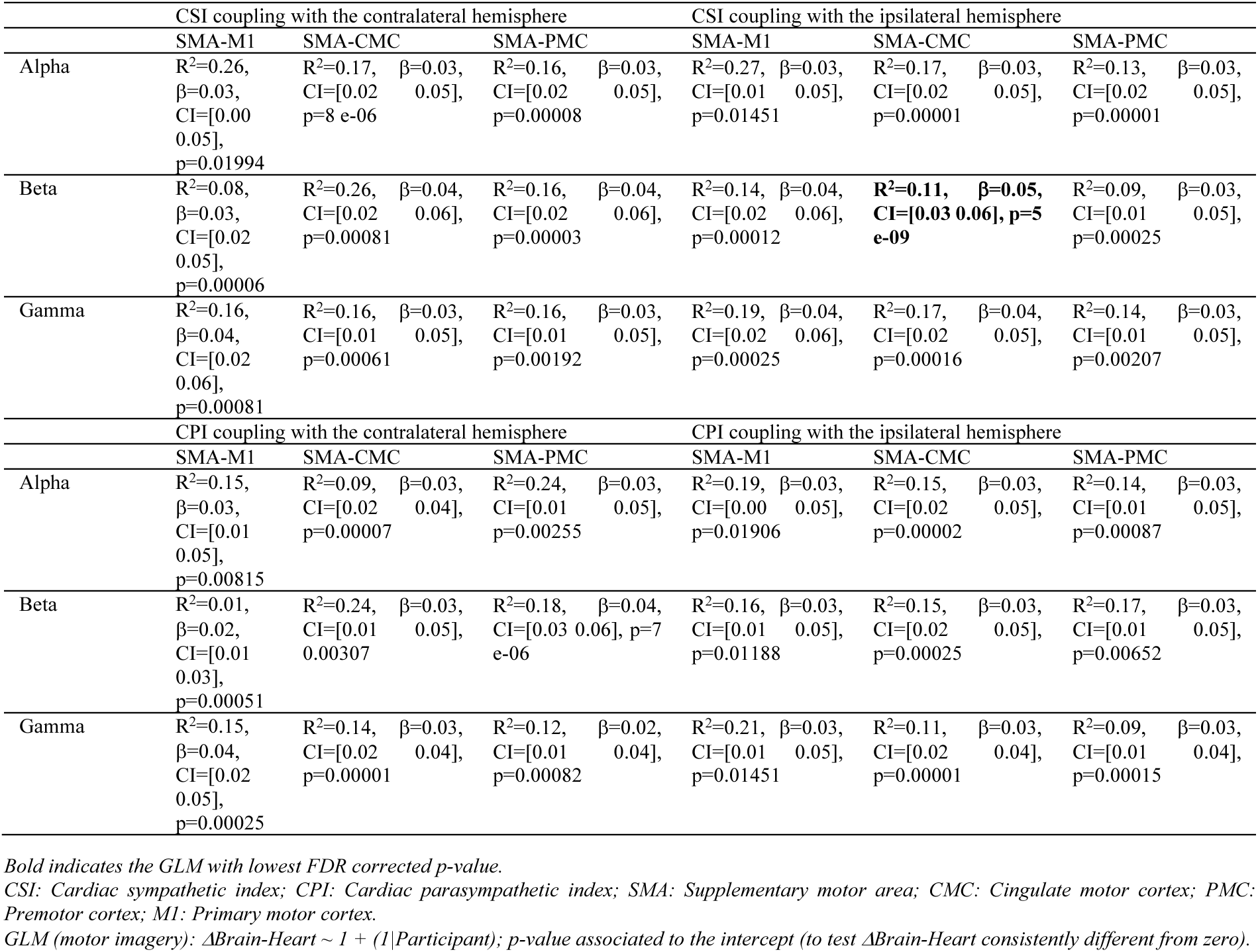
Generalized Linear Models (GLMs) of the change of Heart-Cortico-Cortical relationships in motor imagery, with respect to resting state (11Brain-Heart). This table presents p-values from GLMs assessing CSI-SMA-other motor area interactions during motor imagery. Separate models were fitted to identify the triads consistently modulated by motor imagery.

Figure 2 illustrates changes in the coupling between the CSI and ipsilateral brain connectivity from the SMA to the CMC in the beta band. A paired Mann-Whitney test comparing each motor imagery run to its corresponding resting state (occurring at the beginning of the session) showed significant modulation of brain-heart coupling in most runs across all sessions. As expected, brain-heart coupling decreased during motor imagery compared to resting state. Importantly, this effect was not sustained by brain connectivity from the SMA to the CMC. A decrease in CSI was observed in most runs, but only during the last two sessions, which was mostly triggered by a modulation on heart rate (see Supplementary Figure 1), but these changes in CSI were not correlated with changes in the thalamus (see Supplementary Figure 2). While this decrease was consistent across most of the participants, its amplitude remained low. In addition, we controlled for those effects in the contralateral hemisphere (see Supplementary Figure 3). We found that brain-heart coupling was also modulated in the contralateral hemisphere during motor imagery, but this change occurred in fewer runs compared to the ipsilateral hemisphere.

**Figure 2.**
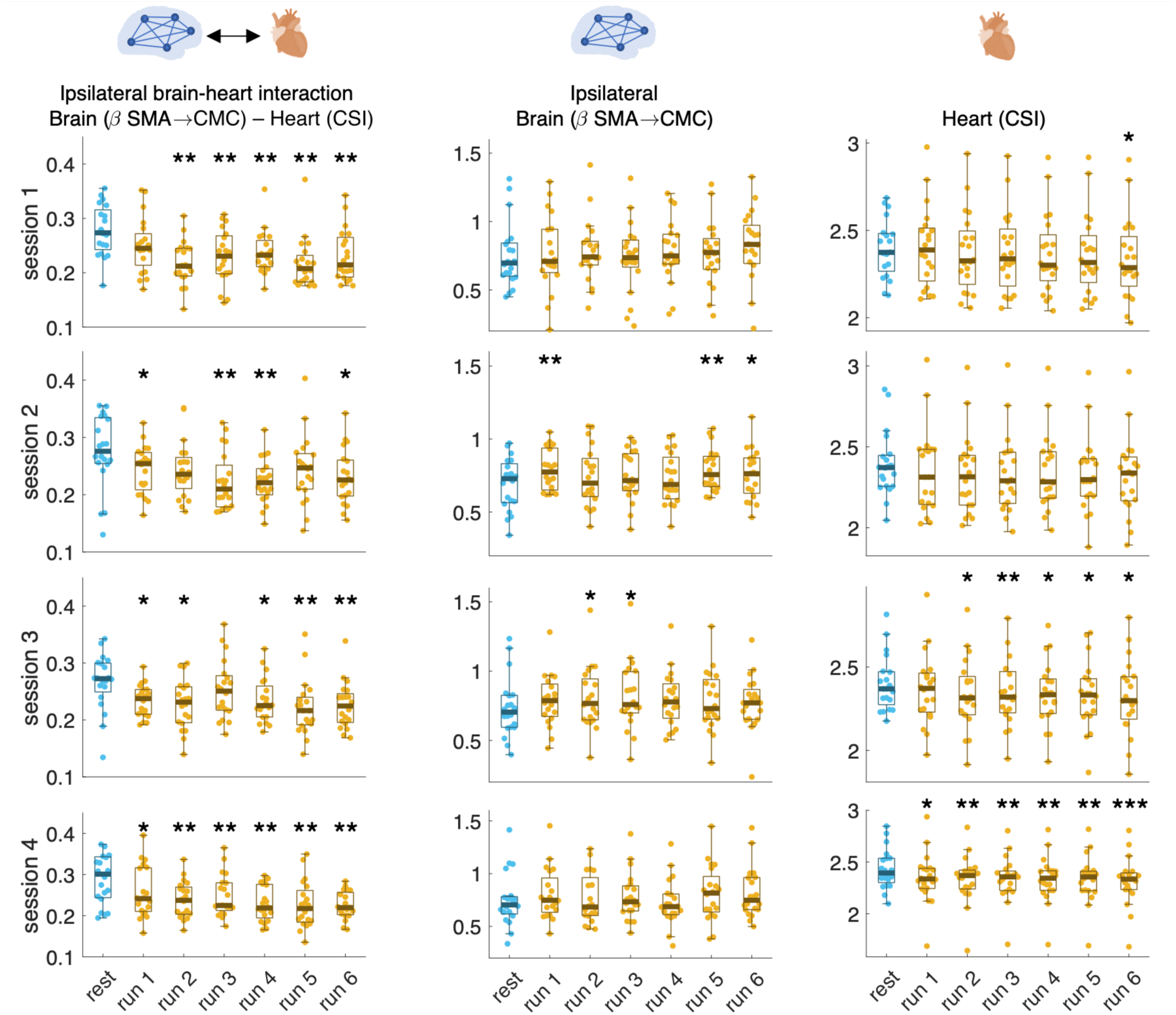
Brain-heart coupling changes during motor imagery. Changes are shown across the short term (runs, horizontal axis) and long term (sessions, vertical axis). Each data point represents one participant (n = 20). Blue markers indicate the resting-state measurements at the beginning of each session, while yellow markers represent the motor imagery runs. The first column shows the coupling between the Cardiac Sympathetic Index (CSI) and the directed connectivity from the Supplementary Motor Area (SMA) to the Cingulate Motor Cortex (CMC) in the ipsilateral hemisphere, within the beta frequency band. This brain-heart triad was identified as the most robust in capturing significant changes from rest across all sessions and runs, as determined by a Generalized Linear Model (see Table 1). The second column displays changes in the directed connectivity from SMA to CMC (ipsilateral, beta band) alone. The third column presents changes in CSI alone. Non-parametric paired Mann–Whitney tests were used to compare each motor imagery run to the corresponding resting-state baseline. Uncorrected p-values are indicated as follows: * p < 0.05, ** p < 0.01, *** p < 0.001.

We then tested whether the brain-heart coupling modulations observed during motor imagery could be explained by thalamo-cortical dynamics, focusing on the SMA and CMC (see Supplementary Figures 4 and 5). We found that bidirectional beta connectivity between the thalamus and SMA, as well as connectivity from CMC to the thalamus, were primarily modulated by motor imagery. However, these modulations occurred in fewer motor imagery runs compared to the coupling between CSI and connectivity from SMA to CMC.

Supplementary analyses confirmed that MEG-EMG coherence was minimal in the 30-100 Hz range and that cardiac indices were not systematically related to breathing rates (Supplementary Fig. 13 and 14). These results indicate that the primary effects are unlikely to be driven by EMG or respiration-related confounds.

### The link between motor imagery accuracy and brain-heart interactions

We investigated whether physiological dynamics could predict fluctuations in motor imagery accuracy while accounting for individual differences across participants and sessions. Using a GLM, we tested whether changes in brain-heart interaction from rest to motor imagery could predict accuracy for the previously identified brain-heart interaction triads. As shown in Table 2, accuracy was best predicted by changes in brain-heart coupling between the CSI and the directed connectivity from the ipsilateral SMA to the PMC in the alpha band, and by the brain-heart coupling between the CPI and the directed connectivity from the ipsilateral SMA to the M1 in the gamma band

**Table 2.** Generalized Linear Models (GLMs) of the change of Heart-Cortico-Cortical relationships in motor imagery, with respect to resting state (11Brain-Heart). This table presents p-values from GLMs assessing CSI-SMA-other motor area interactions during motor imagery. Separate models were fitted to identify the triads associated with general accuracy.

|  | CSI coupling with the contralateral hemisphere |  |  | CSI coupling with the ipsilateral hemisphere |  |  |
| --- | --- | --- | --- | --- | --- | --- |
|  | SMA-M1 | SMA-CMC | SMA-PMC | SMA-M1 | SMA-CMC | SMA-PMC |
| Alpha | $R^2=0.37$, $\beta=10.28$,<br>CI=[-3.85 24.42],<br>$p=0.33755$ | $R^2=0.38$, $\beta=-15.73$,<br>CI=[-31.89 0.43],<br>$p=0.22535$ | $R^2=0.37$, $\beta=-8.21$,<br>CI=[-23.15 6.72],<br>$p=0.53922$ | $R^2=0.37$, $\beta=-10.63$,<br>CI=[-24.54 3.28],<br>$p=0.33755$ | $R^2=0.37$, $\beta=-14.55$,<br>CI=[-31.42 2.33],<br>$p=0.29775$ | **$R^2=0.39$, $\beta=-25.32$,**<br>**CI=[-39.89 -10.76],**<br>**$p=0.01579$** |
| Beta | $R^2=0.37$, $\beta=-2.83$,<br>CI=[-15.50 9.85],<br>$p=0.79145$ | $R^2=0.37$, $\beta=3.41$,<br>CI=[-12.92 19.74],<br>$p=0.79145$ | $R^2=0.37$, $\beta=-2.47$,<br>CI=[-15.88 10.93],<br>$p=0.80669$ | $R^2=0.37$, $\beta=6.57$,<br>CI=[-5.48 18.63],<br>$p=0.53922$ | $R^2=0.37$, $\beta=-2.66$,<br>CI=[-18.92 13.61],<br>$p=0.81631$ | $R^2=0.37$, $\beta=3.26$,<br>CI=[-10.35 16.88],<br>$p=0.79145$ |
| Gamma | $R^2=0.37$, $\beta=-6.74$,<br>CI=[-19.51 6.03],<br>$p=0.54041$ | $R^2=0.37$, $\beta=-4.16$,<br>CI=[-20.60 12.29],<br>$p=0.79145$ | $R^2=0.37$, $\beta=-5.00$,<br>CI=[-19.18 9.17],<br>$p=0.70346$ | $R^2=0.37$, $\beta=-5.87$,<br>CI=[-19.08 7.35],<br>$p=0.59849$ | $R^2=0.37$, $\beta=-11.66$,<br>CI=[-27.91 4.60],<br>$p=0.33755$ | $R^2=0.38$, $\beta=15.85$,<br>CI=[2.05 29.66],<br>$p=0.13005$ |
|  | CPI coupling with the contralateral hemisphere |  |  | CPI coupling with the ipsilateral hemisphere |  |  |
|  | SMA-M1 | SMA-CMC | SMA-PMC | SMA-M1 | SMA-CMC | SMA-PMC |
| Alpha | $R^2=0.41$, $\beta=-14.28$,<br>CI=[-47.26 18.71],<br>$p=0.59849$ | $R^2=0.41$, $\beta=-27.79$,<br>CI=[-70.80 15.22],<br>$p=0.83003$ | $R^2=0.41$, $\beta=-26.23$,<br>CI=[-60.12 7.67],<br>$p=0.95015$ | $R^2=0.41$, $\beta=6.26$,<br>CI=[-26.35 38.88],<br>$p=0.58174$ | $R^2=0.41$, $\beta=-6.94$,<br>CI=[-48.59 34.72],<br>$p=0.93696$ | $R^2=0.42$, $\beta=-30.15$,<br>CI=[-62.80 2.51],<br>$p=0.79145$ |
| Beta | R<sup>2</sup>=0.41, $\beta$=2.46, CI=[-30.97 35.90], p=0.33755 | R<sup>2</sup>=0.41, $\beta$=-0.58, CI=[-38.22 37.05], p=0.33755 | R<sup>2</sup>=0.42, $\beta$=-22.33, CI=[-54.20 9.54], p=0.33755 | R<sup>2</sup>=0.41, $\beta$=-9.89, CI=[-38.05 18.28], p=0.75396 | R<sup>2</sup>=0.42, $\beta$=-23.50, CI=[-64.18 17.18], p=0.13005 | R<sup>2</sup>=0.41, $\beta$=8.68, CI=[-26.63 43.99], p=0.58174 |
| Gamma | R<sup>2</sup>=0.41, $\beta$=10.57, CI=[-22.75 43.88], p=0.27390 | R<sup>2</sup>=0.41, $\beta$=9.67, CI=[-27.59 46.93], p=0.13005 | R<sup>2</sup>=0.41, $\beta$=15.11, CI=[-15.74 45.97], p=0.13005 | **R<sup>2</sup>=0.39, $\beta$=29.17, CI=[2.68 55.67], p=0.01579** | R<sup>2</sup>=0.41, $\beta$=-6.96, CI=[-45.97 32.05], p=0.22535 | R<sup>2</sup>=0.41, $\beta$=-6.82, CI=[-37.99 24.36], p=0.13005 |

*Bold indicates the GLM with lowest FDR corrected p-value.*
*CSI: Cardiac sympathetic index; CPI: Cardiac parasympathetic index; SMA: Supplementary motor area; CMC: Cingulate motor cortex; PMC: Premotor cortex; M1: Primary motor cortex.*
*GLM (motor imagery accuracy): Accuracy ~ $\Delta$ Brain-Heart + (1|Participant); p-value associated to $\Delta$ Brain-Heart*

Figure 3 shows a scatter plot examining the relationship between motor imagery accuracy and the brain-heart coupling between the CSI and the directed connectivity from the ipsilateral SMA to the PMC in the alpha band, and by the brain-heart coupling between the CPI and the directed connectivity from the ipsilateral SMA to the M1 in the gamma band. A Spearman correlation and repeated measures Spearman correlation analysis across all data points (n=480) revealed a significant correlation between changes in brain-heart coupling (relative to rest) and accuracy in the corresponding run, only in the case of gamma SMA-M1-CPI coupling. Greater changes in brain-heart coupling from resting state were associated with higher accuracy. This effect was not replicated by testing the correlation in brain connectivity and CSI separately, or when testing the brain-heart interaction in the contralateral hemisphere (see Supplementary Figure 6)

**Figure 3.**
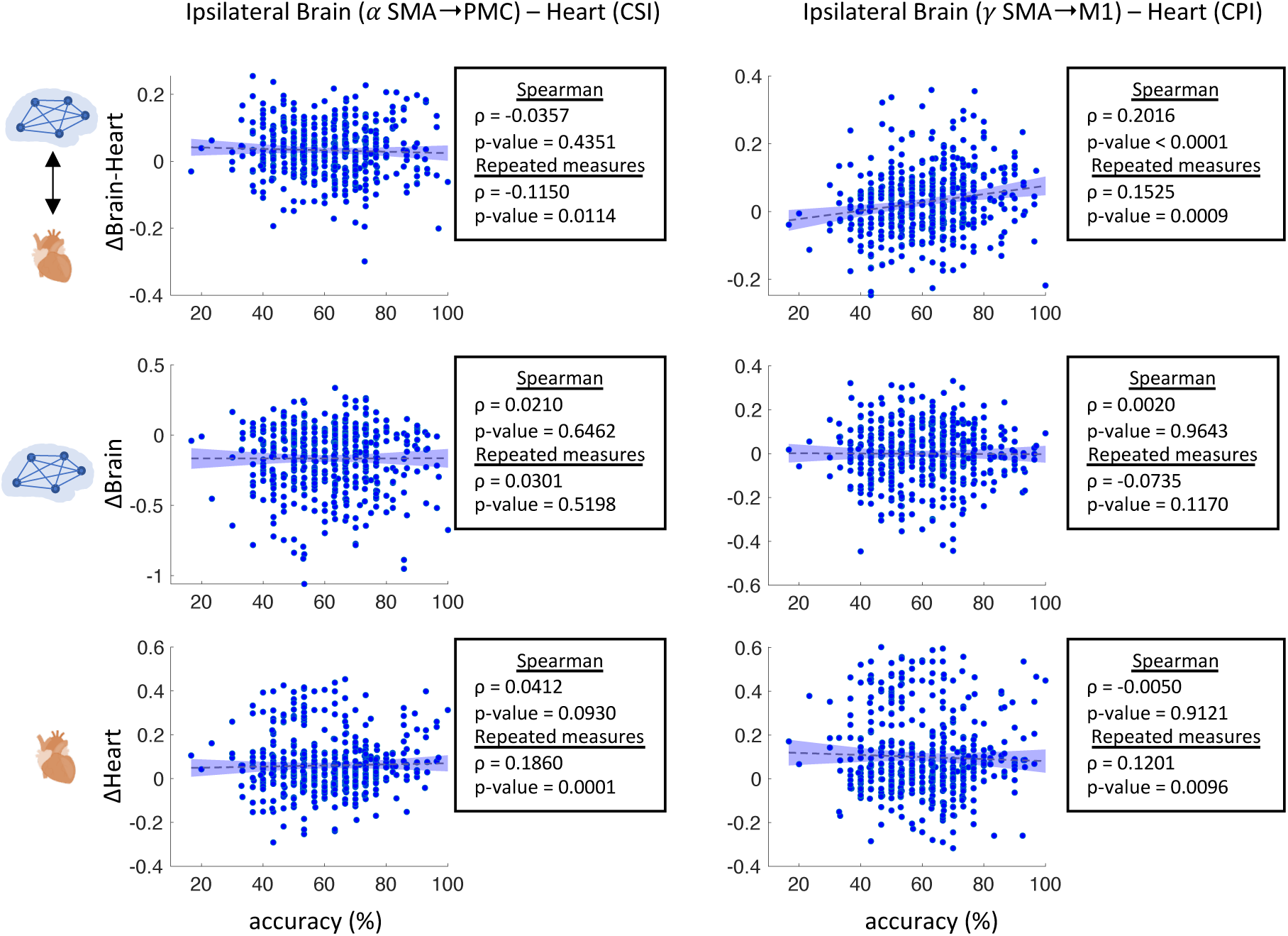
Relationship between the changes in brain-heart coupling and motor imagery accuracy. Each data point represents one participant (n = 20), in one specific run. Blue markers indicate the difference between the resting-state and motor imagery run. The top panel shows the coupling between the Cardiac Sympathetic or Parasympathetic Index (CSI or CPI) and the directed connectivity from the Supplementary Motor Area (SMA) to the Premotor Cortex (PMC) or the primary motor cortex (M1), both in the ipsilateral hemisphere, within the alpha and gamma frequency bands. The middle panel displays changes in the directed connectivity from SMA to PMC and SMA to M1, alone. The third column presents changes in CSI and CPI alone. Non-parametric Spearman correlation tests were used to evaluate the relationship between the change in brain-heart coupling and accuracy

We then examined whether the relationship between brain-heart coupling and motor imagery accuracy could be accounted for by changes in thalamo-cortical connectivity involving the SMA and PMC (see Supplementary Figures 7-9). We found a negative correlation with directed gamma connectivity from the SMA to the thalamus in the contralateral hemisphere, and a positive correlation from the thalamus to the SMA in the contralateral hemisphere. As with the correlations between brain-heart coupling and motor imagery accuracy, the correlation coefficients were relatively low.

### Cardiac-cortico-cerebellar interactions shape longitudinal motor imagery learning

We confirmed that group-level mean motor imagery accuracy consistently improved across sessions. This increase was sustained over time, leading to a significant difference by the final session, indicating an underlying a learning effect across sessions (Mann-Whitney test on the mean accuracy per session, Session 4 > Session 1, p = 0.0006), with a stronger effect in session 3 to 4 (Mann-Whitney test on the mean accuracy per session, Session 2 > Session 1, p = 0.2574; Session 3 > Session 2, p = 0.0793; Session 4 > Session 3, p = 0.0010). To further investigate cortico-cerebellar dynamics associated with motor imagery learning, we examined theta band connectivity patterns from the Cerebellar Crus I to the SMA, PMC and M1, focusing on their relationship with cardiac sympathetic and parasympathetic activity (see Methods for the selection criteria of brain regions and frequency bands). Using GLMs (Table 3) to identify the triad that best captured motor imagery learning dynamics, we found that brain-heart coupling between CPI and directed connectivity from the Cerebellar Crus I to the contralateral SMA in the theta band significantly reflected changes in the relationship between brain-heart coupling and accuracy across sessions.

**Table 3.** Generalized Linear Models (GLMs) of the change of Heart-Cerebellar-Cortical relationships in motor imagery learning, with respect to resting state (11Brain-Heart). This table presents the GLMs assessing CSI-Cerebellar Crus I-other motor area interactions during motor imagery. The models were fitted to identify the triads associated with the longer-term changes in motor imagery accuracy across sessions, specifically the interaction effect (11Brain-Heart ξ Session).

|  |  | Cerebellar Crus-contralateral hemisphere |  |  | Cerebellar Crus-ipsilateral hemisphere |  |  |
| --- | --- | --- | --- | --- | --- | --- | --- |
|  |  | M1 | PMC | SMA | M1 | PMC | SMA |
| CSI | R<sup>2</sup> | R<sup>2</sup>=0.49 | R<sup>2</sup>=0.49 | R<sup>2</sup>=0.49 | R<sup>2</sup>=0.49 | R<sup>2</sup>=0.49 | R<sup>2</sup>=0.49 |
| | *Brain-Heart × Session effect* | $\beta=-1.38$, CI=[-12.02 9.26], $p=0.79911$ | $\beta=5.07$, CI=[-7.61 17.76], $p=0.64854$ | $\beta=10.70$, CI=[-1.61 23.01], $p=0.32717$ | $\beta=7.98$, CI=[-1.79 17.75], $p=0.32717$ | $\beta=-5.60$, CI=[-17.48 6.27], $p=0.60717$ | $\beta=3.42$, CI=[-10.39 17.22], $p=0.68382$ |
| | *ΔBrain-Heart effect* | $\beta=-0.20$, CI=[-29.86 29.47], $p=0.98962$ | $\beta=-24.45$, CI=[-60.20 11.31], $p=0.45704$ | $\beta=-25.00$, CI=[-58.37 8.36], $p=0.45704$ | $\beta=-18.73$, CI=[-46.80 9.34], $p=0.45704$ | $\beta=15.75$, CI=[-17.53 49.02], $p=0.60505$ | $\beta=-14.43$, CI=[-50.13 21.27], $p=0.64099$ |
| | *Session effect* | $\beta=4.40$, CI=[3.47 5.34], $p<0.0001$ | $\beta=4.26$, CI=[3.34 5.18], $p<0.0001$ | $\beta=3.94$, CI=[2.98 4.90], $p<0.0001$ | $\beta=4.13$, CI=[3.25 5.01], $p<0.0001$ | $\beta=4.49$, CI=[3.60 5.37], $p<0.0001$ | $\beta=4.21$, CI=[3.28 5.15], $p<0.0001$ |
| CPI | R<sup>2</sup> | R<sup>2</sup>=0.49 | R<sup>2</sup>=0.49 | R<sup>2</sup>=0.50 | R<sup>2</sup>=0.50 | R<sup>2</sup>=0.50 | R<sup>2</sup>=0.49 |
| | *Brain-Heart × Session effect* | $\beta=3.00$, CI=[-8.22 14.22], $p=0.68382$ | $\beta=8.07$, CI=[-6.00 22.14], $p=0.52048$ | **$\beta=17.38$, CI=[5.60 29.16], $p=0.04685$** | $\beta=2.58$, CI=[-7.84 13.00], $p=0.68382$ | $\beta=15.71$, CI=[3.63 27.78], $p=0.06536$ | $\beta=8.84$, CI=[-4.66 22.35], $p=0.47738$ |
| | *ΔBrain-Heart effect* | $\beta=2.30$, CI=[-28.93 33.54], $p=0.96535$ | $\beta=-11.09$, CI=[-51.04 28.86], $p=0.77420$ | $\beta=-45.16$, CI=[-79.65 -10.66], $p=0.12488$ | $\beta=6.42$, CI=[-20.96 33.80], $p=0.77420$ | $\beta=-34.41$, CI=[-65.85 -2.96], $p=0.19226$ | $\beta=-19.10$, CI=[-57.12 18.93], $p=0.60505$ |
| | *Session effect* | $\beta=4.22$, CI=[3.31 5.13], $p<0.0001$ | $\beta=4.13$, CI=[3.24 5.01], $p<0.0001$ | $\beta=3.63$, CI=[2.66 4.60], $p<0.0001$ | $\beta=4.33$, CI=[3.40 5.26], $p<0.0001$ | $\beta=3.92$, CI=[3.01 4.82], $p<0.0001$ | $\beta=4.04$, CI=[3.05 5.03], $p<0.0001$ |

*Bold indicates the GLM with lowest FDR corrected p-value in the interaction effect ( $\Delta$ Brain-Heart × Session).*
CSI: Cardiac sympathetic index; CPI: Cardiac parasympathetic index; SMA: Supplementary motor area; CMC: Cingulate motor cortex; PMC: Premotor cortex; M1: Primary motor cortex
GLM: Accuracy ∼ Session + $\Delta$ Brain-Heart + $\Delta$ Brain-Heart × Session + (1|Participant)

**Table 4.**
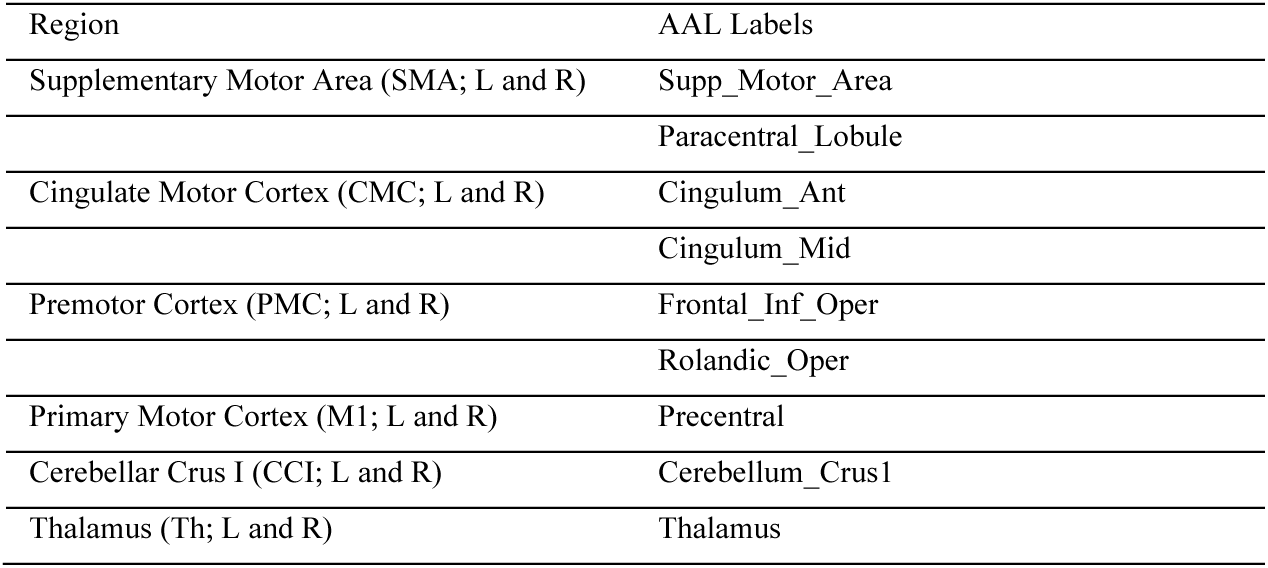
Regions of interest.

To corroborate these findings, we averaged accuracy across all runs per session for each participant to examine whether the correlation between accuracy and brain-heart coupling (CPI–Cerebellar Crus I–SMA in the theta band) improved over sessions. As shown in Figure 4, the Spearman correlation coefficient increased in the third session, reaching a very strong correlation in the fourth session. Furthermore, we found that this effect was stronger when considering the connectivity from the right Cerebellar Crus I to the left SMA (see Supplementary Figure 10). This effect was not observed when controlling for changes in brain connectivity and CPI separately. These results suggest that while Cardiac–Motor cortex connectivity is linked to motor imagery accuracy, the parasympathetic nervous system and the cerebellum may play an active role in the neuroplasticity underlying longitudinal motor imagery learning.

**Figure 4.**
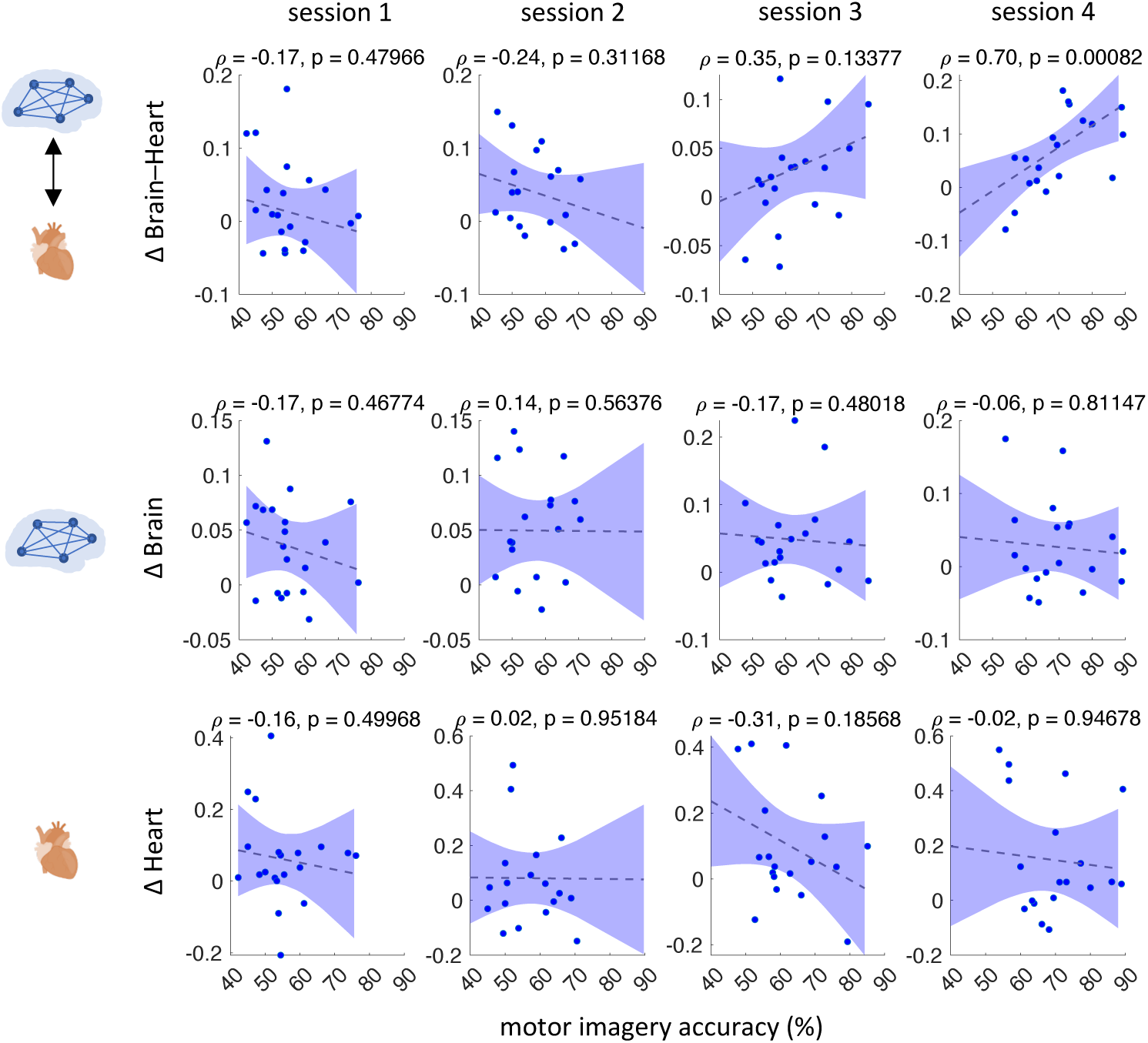
Relationship between the changes in brain-heart coupling and longitudinal motor imagery learning. Each data point represents one participant (n = 20), in one specific session. Blue markers indicate the mean difference between the resting-state and motor imagery session. The top panel shows the coupling between the Cardiac parasympathetic Index (CPI) and the directed connectivity from the Cerebellar Crus I to the Supplementary Motor Area (SMA) in the contralateral hemisphere, within the theta frequency band. This brain-heart triad was identified as the most robust in capturing changes in accuracy associated to longer-term learning across sessions, as determined by a Generalized Linear Model (see Table 2). The middle panel displays changes in the directed connectivity from Cerebellar Crus I to SMA (contralateral, theta band) alone. The third column presents changes in CPI alone.

Finally, we investigated whether the thalamus was involved in the cortico-cerebellar-cardiac relationship associated with motor imagery learning (see Supplementary Figures 11 and 12). We found that the progressive changes in correlations with motor imagery accuracy—indicative of underlying motor imagery learning—were not present when analyzing directed connectivity from the cerebellum to the thalamus or from the thalamus to the SMA.

## Discussion

Exploring brain oscillations during motor imagery could enhance our understanding of its physiological basis. However, there is limited evidence connecting widespread interactions between brain and peripheral neural dynamics. Neural dynamics involve complex processes across brain networks, often studied in isolation from other organs. This gap extends to neurotechnology development: while some efforts have included cardiac information in brain-computer interfaces, these attempts mostly add extra features without quantifying the functional interactions between brain and heart (Fló et al., 2025; Georgaras and and Vourvopoulos, 2025; Kaufmann et al., 2012; Pfurtscheller et al., 2013; Scherer et al., 2007). Recent research shows that cortical functional interactions significantly improve the assessment of motor imagery performance (Cattai et al., 2021; Corsi et al., 2020; Venot et al., 2024). Moreover, the viscera actively responds to and shapes different cognitive and sensorimotor contexts, indicating that organs like the heart act as an active influencer in neural activity (Azzalini et al., 2019; Candia-Rivera, 2022; Engelen et al., 2023; Silvani et al., 2016). In this study, we aimed to demonstrate that cortical connectivity patterns are linked to cardiac dynamics, and that the change of this association may provide more information than considering brain or heart biomarker amplitudes alone. We showed that motor imagery is associated with a dynamic neural segregation, as shown in functional disconnections between CSI and directed connectivity from the SMA to other motor regions. This effect was also found in the coupling between CPI and directed connectivity from the cerebellum to the SMA, in this case signaling longitudinal motor imagery learning. These results suggest that mechanisms operating within the brain-heart axis serve as biomarkers of motor imagery, but also to track learning changes.

Motor imagery engages a well-documented network of brain regions involved in motor planning and execution (Cattai et al., 2021; Corsi et al., 2020; Venot et al., 2024; Virameteekul and Bhidayasiri, 2022). The motor imagery network presents significant overlap with those involved in motor preparation (Jeannerod, 1994) and actual motor execution (Hétu et al., 2013). For instance, premotor cortices participate in schema instantiation (Jeannerod, 1994). The SMA is consistently activated during motor imagery (Picard and Strick, 2003), reflecting its role in motor planning even in the absence of actual movement. In contrast, the M1 is typically associated with motor execution, though some studies report its activation during motor imagery as well (Porro et al., 1996). However, the prevalence of SMA activation across studies supports its crucial role in the covert rehearsal of movements (Roland et al., 1980). In our results, we find that CSI and SMA connectivity with the CMC are specifically engaged during motor imagery, distinguishing this state from resting and further linking SMA-PMC and SMA-M1 interactions to motor imagery accuracy. Previous research has demonstrated the active engagement of SMA and CMC in the earliest stages of pre-movement activity (Cunnington et al., 2003; Nguyen et al., 2014). This suggests that SMA serves as a core hub in the motor imagery process, integrating planning-related activity with motor execution areas. Importantly, cingulate cortices are considered integral hubs of the central autonomic network (Beissner et al., 2013; Valenza et al., 2019). Additionally, the cerebellum, widely recognized for its role in motor learning (Welniarz et al., 2021), in our results also plays a key role in longitudinal learning (a few days apart) in motor imagery training, reinforcing its function in refining motor representations over time.

Motor-sympathetic neural circuits are activated during various physiological and emotionally motivated behaviors (Dum et al., 2019). These circuits synthesize serotonin or contain neuroactive peptides like vasopressin, oxytocin, orexins, and melanin-concentrating hormone(Kerman, 2008). Still, standard markers of heart rate and heart rate variability do not always show relationships with motor imagery performance or ability(Peixoto Pinto et al., 2017). Anticipatory cardiac deceleration, rather than cardiac acceleration, often followed the stimulus, suggesting that the motoric inhibition required during response selection leads to a phasic cardiac deceleration (Jennings et al., 1991). The regulation of skeletal muscle activity is closely linked to sympathetic nervous system activity (Mitchell and Victor, 1996). Notably, research has suggested that cardiac sympathetic activity may serve as a more effective noninvasive marker of muscle neurotransmitter dynamics than skin conductance (Valenza et al., 2020, p. 0; Wallin and Fagius, 1988). For this, skin conductance monitoring in motor-related tasks may not outperform those from the heart, in cases such as hybrid brain-computer interfaces. Importantly, changes in heart and respiration rates can be found as early as participants start to perform motor imagery (Oishi et al., 2000; Oishi and Maeshima, 2004). Indeed, sympathetic tone may change in preparation of movement (Critchley, 2002) and handgrip(Leuenberger et al., 2003). Still, sympathetic markers alone do not seem to suffice as input features in brain-computer interfaces, as these approaches have not been widely employed (Choi et al., 2017).

The role of cardiac parasympathetic activity in motor imagery remains unclear, as fewer studies have examined changes in parasympathetic markers. However, baseline vagal tone has been linked to motor imagery ability and vividness(Sebastiani et al., 2019). More research on the parasympathetic nervous system focuses on its role in bodily self-awareness (Candia-Rivera et al., 2024a). Recent evidence showed that interventions that enhance the anchoring of the self to the body, such as social touch, engage an increase in cardiac parasympathetic activity (Candia-Rivera et al., 2025a). Moreover, those cardiac dynamics covary with the functional activation of frontoparietal networks in alpha and gamma bands (Candia-Rivera et al., 2025b). This implies a direct relationship between the parasympathetic nervous system and the bodily self, likely overlapping with those mechanisms necessary for motor imagery and sensorimotor integration (Evans and Blanke, 2013; Serino et al., 2022).

Brain-heart interactions are mediated, in part, by sympathetic and parasympathetic nervous systems (Chen et al., 2021), where afferent signals from the heart relate to brain activity in areas involved in perceptual awareness and sensorimotor functions (Engelen et al., 2023). While the brain dynamics of motor imagery have been extensively studied at the brain and autonomic levels, separately, their interactions potentially captured through brain-heart couplings remained highly uncharted. Brain-heart interactions are part of interoception (Chen et al., 2021)—the monitoring, interpretation and regulation of internal signals like heartbeat. And interoception has been closely attached to bodily self-awareness—the sensations of being a body and having control over it (Candia-Rivera et al., 2024a). In the motor-related context, research showed that the phase of the cardiac cycle during stimulus presentation can influence action execution and response times (Al et al., 2023; Palser et al., 2021). However, evidence on brain-heart interactions during motor imagery remains limited, particularly regarding oscillatory interactions. Existing studies have mainly focused on the cardiac cycle but not on heart rate fluctuations. Particularly, the cardiac phase appears to influence motor imagery performance, with greater suppression of alpha and beta activity during diastole(Lai et al., 2024).

We found a strong relationship between a cardiac-cortico-cerebellar circuit and motor imagery learning. Consistent with this interpretation, most participants exhibited an increase in motor imagery accuracy from the first to the fourth session. However, this improvement was not linear, as accuracy gains were not sustained on a session-by-session basis, suggesting fluctuating learning dynamics rather than a monotonic progression. As expected, the cerebellum appears as a key node in general motor learning, in particular the cerebellar crus (Gao et al., 2018; Kipping et al., 2013; Stoodley et al., 2012). Noteworthy, the cerebellum is among the brain regions associated with the central autonomic network because of its correlates with cardiac autonomic dynamics (Valenza et al., 2024). The specific relationship in our results was with respect to cardiac parasympathetic dynamics, whose link with the cerebellum occurs through vagal inputs via the nucleus tractus solitarius (Andresen and Kunze, 1994). This non-linear behavioral improvement may reflect alternating phases of exploration, consolidation, and adaptation during motor imagery learning, and aligns with the notion that cerebellar mechanisms support error prediction and updating rather than continuous performance gains (Dayan and Cohen, 2011; Wolpert et al., 1998). Our results suggest that the cerebellum, while involved in motor imagery learning, also integrates cardiac inputs during this process. This points to a potential feedback loop between motor imagery, the parasympathetic nervous system, and error prediction mechanisms.

During motor-related tasks, cortical changes are typically lateralized in M1. In motor imagery, with mixed results (Corsi et al., 2020; Nam et al., 2011; Porro et al., 2000), the lateralization is observed as well in M1 and PMC. Instead, ipsilateral and bilaterial activity is found in SMA, cerebellum, frontoparietal networks and other frontal regions (Decety, 1996). In our results, the segregation from cardiac dynamics was primarily observed in the ipsilateral SMA during all motor imagery runs, and in the contralateral SMA in those dynamics associated with motor imagery learning. Taken together, these findings suggest that lateralization may reflect task-specific functions and depend on the biomarker under investigation. Importantly, since our analysis focuses on changes in oscillatory couplings, rather than on absolute increases or decreases in physiological activity, our results should not be directly compared to studies examining event-related potentials or event-related (de)synchronizations within specific frequency bands (Lu and Yin, 2015).

It is important to note that a reduction in statistical dependence, as captured by the MIC, does not necessarily indicate a loss of interaction but may reflect a transition between interaction regimes or changes in the dominant physiological processes involved. In the present context, such apparent uncoupling may index a reorganization of brain-heart interactions during motor imagery rather than disengagement or causal suppression. Importantly, the present findings should be interpreted as associative, and causal inferences will require future studies incorporating perturbation-based approaches or explicit causal modeling frameworks.

Inter-individual variability is a hallmark of motor imagery and BCI paradigms (Vidaurre and Blankertz, 2010), with a substantial proportion of participants often classified as non-responders or illiterates. Rather than treating this variability as a limitation, the present study explicitly aims to investigate physiological factors underlying variability in performance across individuals.

Our study has certain limitations, including the use of source-reconstructed brain signals from structures such as the thalamus and the cerebellum, as well as a modest sample size that limits the generalizability of the findings, particularly for subgroup analyses. In particular, reconstruction of deep brain structures with MEG remains challenging: although individual MRIs and a conservative inverse solution were used to mitigate localization uncertainty, MEG sensitivity decreases with source depth, and precise localization of subcortical sources cannot be guaranteed. For this reason, cerebellar analyses were restricted a priori to Crus I, and thalamic results were treated as secondary. Accordingly, our interpretations focus on relative interaction patterns and group-level effects rather than absolute source amplitudes or precise anatomical localization.

Nevertheless, it represents the first attempt to provide mechanistic insights into the interplay between brain and cardiac signals during motor imagery. While similar efforts have been made in the context of hybrid brain–computer interface systems, the underlying mechanisms were often obscured by black-box machine learning approaches. In contrast, the present work aims to elucidate physiologically interpretable interactions between central and peripheral signals. Although sex-related differences in neural and autonomic function are well documented, the present study was not powered to reliably assess sex effects, and no sex-stratified analyses were performed to avoid overinterpretation. We believe our findings can pave the way for more sophisticated frameworks that integrate peripheral neural dynamics not merely as additional features, but as meaningful components of the system’s core design, to be further validated in future studies with larger and more balanced samples.

Taken together, our results indicate that motor imagery is not solely characterized by cortical activation patterns, but by dynamic interactions between cardiac autonomic activity and SMA-driven motor networks. We show that cardiac sympathetic and parasympathetic indices selectively relate to directed connectivity from the SMA to cortical and cerebellar motor-related regions, and that these interactions relate to both imagery accuracy and learning. This converging evidence supports the view that brain-heart interactions constitute a functional dimension of motor imagery, reflecting adaptive engagement and disengagement of motor control circuits rather than static couplings.

## Author contributions

Diego Candia-Rivera: Conceptualization, Methodology, Software, Validation, Formal Analysis, Investigation, Writing Original Draft, Writing Review and Editing.

Mario Chavez: Methodology, Investigation, Supervision, Writing Review and Editing. Fabrizio de Vico Fallani: Methodology, Investigation, Writing Review and Editing.

Marie-Constance Corsi: Data curation, Methodology, Investigation, Supervision, Writing Review and Editing.

## References

Ahn, M., Ahn, S., Hong, J.H., Cho, H., Kim, K., Kim, B.S., Chang, J.W., Jun, S.C., 2013. Gamma band activity associated with BCI performance: simultaneous MEG/EEG study. Front. Hum. Neurosci. 7. 10.3389/fnhum.2013.00848

Akkal, D., Dum, R.P., Strick, P.L., 2007. Supplementary Motor Area and Presupplementary Motor Area: Targets of Basal Ganglia and Cerebellar Output. J. Neurosci. 27, 10659–10673. 10.1523/JNEUROSCI.3134-07.2007

Al, E., Stephani, T., Engelhardt, M., Haegens, S., Villringer, A., Nikulin, V.V., 2023. Cardiac activity impacts cortical motor excitability. PLOS Biology 21, e3002393. 10.1371/journal.pbio.3002393

Al-Wasity, S., Vogt, S., Vuckovic, A., Pollick, F.E., 2021. Upregulation of Supplementary Motor Area Activation with fMRI Neurofeedback during Motor Imagery. eNeuro 8. 10.1523/ENEURO.0377-18.2020

Andersen, L.M., Jerbi, K., Dalal, S.S., 2020. Can EEG and MEG detect signals from the human cerebellum? NeuroImage 215, 116817. 10.1016/j.neuroimage.2020.116817

Andresen, M.C., Kunze, D.L., 1994. Nucleus Tractus Solitarius—Gateway to Neural Circulatory Control. Annual Review of Physiology 56, 93–116. 10.1146/annurev.ph.56.030194.000521

Azzalini, D., Rebollo, I., Tallon-Baudry, C., 2019. Visceral Signals Shape Brain Dynamics and Cognition. Trends in Cognitive Sciences 23, 488–509. 10.1016/j.tics.2019.03.007

Beissner, F., Meissner, K., Bär, K.-J., Napadow, V., 2013. The autonomic brain: an activation likelihood estimation meta-analysis for central processing of autonomic function. J. Neurosci. 33, 10503–10511. 10.1523/JNEUROSCI.1103-13.2013

Berman, B.D., Horovitz, S.G., Venkataraman, G., Hallett, M., 2012. Self-modulation of primary motor cortex activity with motor and motor imagery tasks using real-time fMRI- based neurofeedback. NeuroImage 59, 917–925. 10.1016/j.neuroimage.2011.07.035

Candia-Rivera, D., 2026. Bodily self-consciousness supports motor imagery. Nat Rev Psychol 1–1. 10.1038/s44159-025-00528-9

Candia-Rivera, D., 2022. Brain-heart interactions in the neurobiology of consciousness. Current Research in Neurobiology 3, 100050. 10.1016/j.crneur.2022.100050

Candia-Rivera, D., Boehme, R., Salamone, P.C., 2025a. Autonomic modulations to cardiac dynamics in response to affective touch: Differences between social touch and self-touch. IEEE Transactions on Affective Computing 1–11. 10.1109/TAFFC.2025.3548778

Candia-Rivera, D., Chavez, M., 2025. Cardiac-vagal rhythm echoes on the heartbeat’s mechanosensory imprint in the brain. Commun Biol 8, 1578. 10.1038/s42003-025-08969-x

Candia-Rivera, D., de Vico Fallani, F., Boehme, R., Salamone, P.C., 2025b. Linking heartbeats with the cortical network dynamics involved in self-social touch distinction. Commun Biol 8, 1–13. 10.1038/s42003-024-07448-z

Candia-Rivera, D., de Vico Fallani, F., Chavez, M., 2025c. Robust and time-resolved estimation of cardiac sympathetic and parasympathetic indices. Royal Society Open Science 12, 240750. 10.1098/rsos.240750

Candia-Rivera, D., Engelen, T., Babo-Rebelo, M., Salamone, P.C., 2024a. Interoception, Network Physiology and the Emergence of Bodily Self-Awareness. Neuroscience & Biobehavioral Reviews 165, 105864. 10.1016/j.neubiorev.2024.105864

Candia-Rivera, D., Vidailhet, M., Chavez, M., De Vico Fallani, F., 2024b. A framework for quantifying the coupling between brain connectivity and heartbeat dynamics: Insights into the disrupted network physiology in Parkinson’s disease. Human Brain Mapping 45, e26668. 10.1002/hbm.26668

Catrambone, V., Candia-Rivera, D., Valenza, G., 2024. Intracortical brain-heart interplay: An EEG model source study of sympathovagal changes. Human Brain Mapping 45, e26677. 10.1002/hbm.26677

Cattai, T., Colonnese, S., Corsi, M.-C., Bassett, D.S., Scarano, G., De Vico Fallani, F., 2021. Phase/Amplitude Synchronization of Brain Signals During Motor Imagery BCI Tasks. IEEE Transactions on Neural Systems and Rehabilitation Engineering 29, 1168–1177. 10.1109/TNSRE.2021.3088637

Cavina-Pratesi, C., Monaco, S., Fattori, P., Galletti, C., McAdam, T.D., Quinlan, D.J., Goodale, M.A., Culham, J.C., 2010. Functional Magnetic Resonance Imaging Reveals the Neural Substrates of Arm Transport and Grip Formation in Reach-to-Grasp Actions in Humans. J. Neurosci. 30, 10306–10323. 10.1523/JNEUROSCI.2023-10.2010

Chen, H., Yang, Q., Liao, W., Gong, Q., Shen, S., 2009. Evaluation of the effective connectivity of supplementary motor areas during motor imagery using Granger causality mapping. NeuroImage 47, 1844–1853. 10.1016/j.neuroimage.2009.06.026

Chen, W.G., Schloesser, D., Arensdorf, A.M., Simmons, J.M., Cui, C., Valentino, R., Gnadt, J.W., Nielsen, L., Hillaire-Clarke, C.St., Spruance, V., Horowitz, T.S., Vallejo, Y.F., Langevin, H.M., 2021. The Emerging Science of Interoception: Sensing, Integrating, Interpreting, and Regulating Signals within the Self. Trends in Neurosciences, Special Issue: The Neuroscience of Interoception 44, 3–16. 10.1016/j.tins.2020.10.007

Choi, I., Rhiu, I., Lee, Y., Yun, M.H., Nam, C.S., 2017. A systematic review of hybrid brain-computer interfaces: Taxonomy and usability perspectives. PLOS ONE 12, e0176674. 10.1371/journal.pone.0176674

Cona, G., Marino, G., Semenza, C., 2017. TMS of supplementary motor area (SMA) facilitates mental rotation performance: Evidence for sequence processing in SMA. NeuroImage 146, 770–777. 10.1016/j.neuroimage.2016.10.032

Corsi, M.-C., Chavez, M., Schwartz, D., George, N., Hugueville, L., Kahn, A.E., Dupont, S., Bassett, D.S., De Vico Fallani, F., 2020. Functional disconnection of associative cortical areas predicts performance during BCI training. NeuroImage 209, 116500. 10.1016/j.neuroimage.2019.116500

Critchley, H.D., 2002. Electrodermal responses: what happens in the brain. Neuroscientist 8, 132–142. 10.1177/107385840200800209

Cunnington, R., Windischberger, C., Deecke, L., Moser, E., 2003. The preparation and readiness for voluntary movement: a high-field event-related fMRI study of the Bereitschafts-BOLD response. NeuroImage 20, 404–412. 10.1016/S1053-8119(03)00291-X

Dayan, E., Cohen, L.G., 2011. Neuroplasticity Subserving Motor Skill Learning. Neuron 72, 443–454. 10.1016/j.neuron.2011.10.008

Decety, J., 1996. The neurophysiological basis of motor imagery. Behavioural Brain Research 77, 45–52. 10.1016/0166-4328(95)00225-1

Dekleva, B.M., Chowdhury, R.H., Batista, A.P., Chase, S.M., Yu, B.M., Boninger, M.L., Collinger, J.L., 2024. Motor cortex retains and reorients neural dynamics during motor imagery. Nat Hum Behav 8, 729–742. 10.1038/s41562-023-01804-5

Dum, R.P., Levinthal, D.J., Strick, P.L., 2019. The mind–body problem: Circuits that link the cerebral cortex to the adrenal medulla. Proceedings of the National Academy of Sciences 116, 26321–26328. 10.1073/pnas.1902297116

Dum, R.P., Strick, P.L., 1993. Cingulate Motor Areas, in: Vogt, B.A., Gabriel, M. (Eds.), Neurobiology of Cingulate Cortex and Limbic Thalamus: A Comprehensive Handbook. Birkhäuser, Boston, MA, pp. 415–441. 10.1007/978-1-4899-6704-6_15

Edelman, B.J., Zhang, S., Schalk, G., Brunner, P., Müller-Putz, G., Guan, C., He, B., 2024. Non-invasive Brain-Computer Interfaces: State of the Art and Trends. IEEE Reviews in Biomedical Engineering 1–25. 10.1109/RBME.2024.3449790

Engelen, T., Solcà, M., Tallon-Baudry, C., 2023. Interoceptive rhythms in the brain. Nat Neurosci 26, 1670–1684. 10.1038/s41593-023-01425-1

Evans, N., Blanke, O., 2013. Shared electrophysiology mechanisms of body ownership and motor imagery. NeuroImage 64, 216–228. 10.1016/j.neuroimage.2012.09.027

Facchini, S., Muellbacher, W., Battaglia, F., Boroojerdi, B., Hallett, M., 2002. Focal enhancement of motor cortex excitability during motor imagery: a transcranial magnetic stimulation study. Acta Neurologica Scandinavica 105, 146–151. 10.1034/j.1600-0404.2002.1o004.x

Fischl, B., 2012. FreeSurfer. NeuroImage, 20 YEARS OF fMRI 62, 774–781. 10.1016/j.neuroimage.2012.01.021

Fló, E., Fraiman, D., Sitt, J.D., 2025. Assessing brain-muscle networks during motor imagery to detect covert command-following. BMC Med 23, 68. 10.1186/s12916-025-03846-0

Gao, Z., Davis, C., Thomas, A.M., Economo, M.N., Abrego, A.M., Svoboda, K., De Zeeuw, C.I., Li, N., 2018. A cortico-cerebellar loop for motor planning. Nature 563, 113–116. 10.1038/s41586-018-0633-x

Georgaras, E., and Vourvopoulos, A., 2025. Physiological assessment of brain, cardiovascular, and respiratory changes in multimodal motor imagery brain-computer interface training. Research in Biomedical Engineering and Technology 12, 2471680. 10.1080/29960355.2025.2471680

Gordon, E.M., Chauvin, R.J., Van, A.N., Rajesh, A., Nielsen, A., Newbold, D.J., Lynch, C.J., Seider, N.A., Krimmel, S.R., Scheidter, K.M., Monk, J., Miller, R.L., Metoki, A., Montez, D.F., Zheng, A., Elbau, I., Madison, T., Nishino, T., Myers, M.J., Kaplan, S, Badke D’Andrea, C., Demeter, D.V., Feigelis, M., Ramirez, J.S.B., Xu, T., Barch, D.M., Smyser, C.D., Rogers, C.E., Zimmermann, J., Botteron, K.N., Pruett, J.R., Willie, J.T., Brunner, P., Shimony, J.S., Kay, B.P., Marek, S., Norris, S.A., Gratton, C., Sylvester, C.M., Power, J.D., Liston, C., Greene, D.J., Roland, J.L., Petersen, S.E., Raichle, M.E., Laumann, T.O., Fair, D.A., Dosenbach, N.U.F., 2023. A somato-cognitive action network alternates with effector regions in motor cortex. Nature 617, 351–359. 10.1038/s41586-023-05964-2

Gross, J., Baillet, S., Barnes, G.R., Henson, R.N., Hillebrand, A., Jensen, O., Jerbi, K., Litvak, V., Maess, B., Oostenveld, R., Parkkonen, L., Taylor, J.R., van Wassenhove, V., Wibral, M., Schoffelen, J.-M., 2013. Good practice for conducting and reporting MEG research. NeuroImage 65, 349–363. 10.1016/j.neuroimage.2012.10.001

Grosse-Wentrup, M., Schölkopf, B., 2012. High gamma-power predicts performance in sensorimotor-rhythm brain–computer interfaces. J. Neural Eng. 9, 046001. 10.1088/1741-2560/9/4/046001

Guerra, Z.F., Lucchetti, A.L.G., Lucchetti, G., 2017. Motor Imagery Training After Stroke: A Systematic Review and Meta-analysis of Randomized Controlled Trials. Journal of Neurologic Physical Therapy 41, 205. 10.1097/NPT.0000000000000200

Guger, C., Edlinger, G., Harkam, W., Niedermayer, I., Pfurtscheller, G., 2003. How many people are able to operate an EEG-based brain-computer interface (BCI)? IEEE Transactions on Neural Systems and Rehabilitation Engineering 11, 145–147. 10.1109/TNSRE.2003.814481

Hanakawa, T., Immisch, I., Toma, K., Dimyan, M.A., Van Gelderen, P., Hallett, M., 2003. Functional Properties of Brain Areas Associated With Motor Execution and Imagery. Journal of Neurophysiology 89, 989–1002. 10.1152/jn.00132.2002

Henschke, J.U., Pakan, J.M.P., 2023. Engaging distributed cortical and cerebellar networks through motor execution, observation, and imagery. Front. Syst. Neurosci. 17. 10.3389/fnsys.2023.1165307

Hétu, S., Grégoire, M., Saimpont, A., Coll, M.-P., Eugène, F., Michon, P.-E., Jackson, P.L., 2013. The neural network of motor imagery: An ALE meta-analysis. Neuroscience & Biobehavioral Reviews 37, 930–949. 10.1016/j.neubiorev.2013.03.017

Hoshi, E., Tanji, J., 2007. Distinctions between dorsal and ventral premotor areas: anatomical connectivity and functional properties. Current Opinion in Neurobiology, Cognitive neuroscience 17, 234–242. 10.1016/j.conb.2007.02.003

Jeannerod, M., 1994. The representing brain: Neural correlates of motor intention and imagery. Behavioral and Brain Sciences 17, 187–202. 10.1017/S0140525X00034026

Jennings, J.R., Van Der Molen, M.W., Brock, K., Somsen, R.J.M., 1991. Response Inhibition Initiates Cardiac Deceleration: Evidence from a Sensory-Motor Compatibility Paradigm. Psychophysiology 28, 72–85. 10.1111/j.1469-8986.1991.tb03390.x

Kaufmann, T., Vögele, C., Sütterlin, S., Lukito, S., Kübler, A., 2012. Effects of resting heart rate variability on performance in the P300 brain-computer interface. International Journal of Psychophysiology 83, 336–341. 10.1016/j.ijpsycho.2011.11.018

Kay, S.M., Marple, S.L., 1981. Spectrum analysis—A modern perspective. Proceedings of the IEEE 69, 1380–1419. 10.1109/PROC.1981.12184

Kerman, I.A., 2008. Organization of brain somatomotor-sympathetic circuits. Exp Brain Res 187, 1–16. 10.1007/s00221-008-1337-5

Khan, M.A., Das, R., Iversen, H.K., Puthusserypady, S., 2020. Review on motor imagery based BCI systems for upper limb post-stroke neurorehabilitation: From designing to application. Computers in Biology and Medicine 123, 103843. 10.1016/j.compbiomed.2020.103843

Kilintari, M., Narayana, S., Babajani-Feremi, A., Rezaie, R., Papanicolaou, A.C., 2016. Brain activation profiles during kinesthetic and visual imagery: An fMRI study. Brain Res 1646, 249–261. 10.1016/j.brainres.2016.06.009

Kipping, J.A., Grodd, W., Kumar, V., Taubert, M., Villringer, A., Margulies, D.S., 2013. Overlapping and parallel cerebello-cerebral networks contributing to sensorimotor control: An intrinsic functional connectivity study. NeuroImage 83, 837–848. 10.1016/j.neuroimage.2013.07.027

Lai, G., Landi, D., Vidaurre, C., Bhattacharya, J., Herrojo Ruiz, M., 2024. Cardiac cycle modulates alpha and beta suppression during motor imagery. Cerebral Cortex 34, bhae442. 10.1093/cercor/bhae442

Lebedev, M.A., Nicolelis, M.A.L., 2006. Brain–machine interfaces: past, present and future. Trends in Neurosciences 29, 536–546. 10.1016/j.tins.2006.07.004

Lebon, F., Byblow, W.D., Collet, C., Guillot, A., Stinear, C.M., 2012. The modulation of motor cortex excitability during motor imagery depends on imagery quality. European Journal of Neuroscience 35, 323–331. 10.1111/j.1460-9568.2011.07938.x

Leuenberger, U.A., Mostoufi-Moab, S., Herr, M., Gray, K., Kunselman, A., Sinoway, L.I., 2003. Control of Skin Sympathetic Nerve Activity During Intermittent Static Handgrip Exercise. Circulation 108, 2329–2335. 10.1161/01.CIR.0000093280.40118.30

Lotte, F., Bougrain, L., Cichocki, A., Clerc, M., Congedo, M., Rakotomamonjy, A., Yger, F., 2018. A review of classification algorithms for EEG-based brain–computer interfaces: a 10 year update. J. Neural Eng. 15, 031005. 10.1088/1741-2552/aab2f2

Lotze, M., Erb, M., Flor, H., Huelsmann, E., Godde, B., Grodd, W., 2000. fMRI Evaluation of Somatotopic Representation in Human Primary Motor Cortex. NeuroImage 11, 473–481. 10.1006/nimg.2000.0556

Lu, N., Yin, T., 2015. Motor imagery classification via combinatory decomposition of ERP and ERSP using sparse nonnegative matrix factorization. Journal of Neuroscience Methods 249, 41–49. 10.1016/j.jneumeth.2015.03.031

Makoshi, Z., Kroliczak, Gregory, and van Donkelaar, P., 2011. Human Supplementary Motor Area Contribution to Predictive Motor Planning. Journal of Motor Behavior 43, 303–309. 10.1080/00222895.2011.584085

Malouin, F., Richards, C.L., Jackson, P.L., Dumas, F., Doyon, J., 2003. Brain activations during motor imagery of locomotor-related tasks: A PET study. Human Brain Mapping 19, 47–62. 10.1002/hbm.10103

Mehler, D.M.A., Williams, A.N., Krause, F., Lührs, M., Wise, R.G., Turner, D.L., Linden, D.E.J., Whittaker, J.R., 2019. The BOLD response in primary motor cortex and supplementary motor area during kinesthetic motor imagery based graded fMRI neurofeedback. NeuroImage 184, 36–44. 10.1016/j.neuroimage.2018.09.007

Mitchell, J.H., Victor, R.G., 1996. Neural control of the cardiovascular system: insights from muscle sympathetic nerve recordings in humans. Medicine & Science in Sports & Exercise 28, 60.

Naito, E., Kochiyama, T., Kitada, R., Nakamura, S., Matsumura, M., Yonekura, Y., Sadato, N., 2002. Internally Simulated Movement Sensations during Motor Imagery Activate Cortical Motor Areas and the Cerebellum. J. Neurosci. 22, 3683–3691. 10.1523/JNEUROSCI.22-09-03683.2002

Nam, C.S., Jeon, Y., Kim, Y.-J., Lee, I., Park, K., 2011. Movement imagery-related lateralization of event-related (de)synchronization (ERD/ERS): Motor-imagery duration effects. Clinical Neurophysiology 122, 567–577. 10.1016/j.clinph.2010.08.002

Nguyen, V.T., Breakspear, M., Cunnington, R., 2014. Reciprocal Interactions of the SMA and Cingulate Cortex Sustain Premovement Activity for Voluntary Actions. J. Neurosci. 34, 16397–16407. 10.1523/JNEUROSCI.2571-14.2014

O’Brien, C., Heneghan, C., 2007. A comparison of algorithms for estimation of a respiratory signal from the surface electrocardiogram. Computers in Biology and Medicine 37, 305– 314. 10.1016/j.compbiomed.2006.02.002

Ohata, R., Asai, T., Kadota, H., Shigemasu, H., Ogawa, K., Imamizu, H., 2020. Sense of Agency Beyond Sensorimotor Process: Decoding Self-Other Action Attribution in the Human Brain. Cerebral Cortex 30, 4076–4091. 10.1093/cercor/bhaa028

Oishi, K., Kasai, T., Maeshima, T., 2000. Autonomic Response Specificity during Motor Imagery. Journal of PHYSIOLOGICAL ANTHROPOLOGY and Applied Human Science 19, 255–261. 10.2114/jpa.19.255

Oishi, K., Maeshima, T., 2004. Autonomic Nervous System Activities During Motor Imagery in Elite Athletes. Journal of Clinical Neurophysiology 21, 170.

Oostenveld, R., Fries, P., Maris, E., Schoffelen, J.-M., 2011. FieldTrip: Open Source Software for Advanced Analysis of MEG, EEG, and Invasive Electrophysiological Data. Computational Intelligence and Neuroscience 2011, 9 pages. 10.1155/2011/156869

Owen, A.M., Coleman, M.R., Boly, M., Davis, M.H., Laureys, S., Pickard, J.D., 2006. Detecting awareness in the vegetative state. Science 313, 1402. 10.1126/science.1130197

Palser, E.R., Glass, J., Fotopoulou, A., Kilner, J.M., 2021. Relationship between cardiac cycle and the timing of actions during action execution and observation. Cognition 217, 104907. 10.1016/j.cognition.2021.104907

Pan, J., Tompkins, W.J., 1985. A Real-Time QRS Detection Algorithm. IEEE Transactions on Biomedical Engineering 32, 230–236. 10.1109/TBME.1985.325532

Park, H.-D., Bernasconi, F., Bello-Ruiz, J., Pfeiffer, C., Salomon, R., Blanke, O., 2016. Transient Modulations of Neural Responses to Heartbeats Covary with Bodily Self-Consciousness. J. Neurosci. 36, 8453–8460. 10.1523/JNEUROSCI.0311-16.2016

Pascual-Marqui, R.D., 2002. Standardized low-resolution brain electromagnetic tomography (sLORETA): technical details. Methods Find Exp Clin Pharmacol 24 Suppl D, 5–12.

Peixoto Pinto, T., Mello Russo Ramos, M., Lemos, T., Domingues Vargas, C., Imbiriba, L.A., 2017. Is heart rate variability affected by distinct motor imagery strategies? Physiology & Behavior 177, 189–195. 10.1016/j.physbeh.2017.05.004

Pfurtscheller, G., Solis Escalante, T., Barry, R., Klobassa, D., Neuper, C., Mueller-Putz, G., 2013. Brisk heart rate and EEG changes during execution and withholding of cue-paced foot motor imagery. Frontiers in Human Neuroscience 7.

Picard, N., Strick, P.L., 2003. Activation of the Supplementary Motor Area (SMA) during Performance of Visually Guided Movements. Cerebral Cortex 13, 977–986. 10.1093/cercor/13.9.977

Porro, C.A., Cettolo, V., Francescato, M.P., Baraldi, P., 2000. Ipsilateral involvement of primary motor cortex during motor imagery. European Journal of Neuroscience 12, 3059–3063. 10.1046/j.1460-9568.2000.00182.x

Porro, C.A., Francescato, M.P., Cettolo, V., Diamond, M.E., Baraldi, P., Zuiani, C., Bazzocchi, M., Prampero, P.E. di, 1996. Primary Motor and Sensory Cortex Activation during Motor Performance and Motor Imagery: A Functional Magnetic Resonance Imaging Study. J. Neurosci. 16, 7688–7698. 10.1523/JNEUROSCI.16-23-07688.1996

Rajpura, P., Cecotti, H., Kumar Meena, Y., 2024. Explainable artificial intelligence approaches for brain–computer interfaces: a review and design space. J. Neural Eng. 21, 041003. 10.1088/1741-2552/ad6593

Rebollo, I., Tallon-Baudry, C., 2022. The Sensory and Motor Components of the Cortical Hierarchy Are Coupled to the Rhythm of the Stomach during Rest. J. Neurosci. 42, 2205–2220. 10.1523/JNEUROSCI.1285-21.2021

Reshef, D.N., Reshef, Y.A., Finucane, H.K., Grossman, S.R., McVean, G., Turnbaugh, P.J., Lander, E.S., Mitzenmacher, M., Sabeti, P.C., 2011. Detecting Novel Associations in Large Datasets. Science 334, 1518–1524. 10.1126/science.1205438

Roland, P.E., Larsen, B., Lassen, N.A., Skinhoj, E., 1980. Supplementary motor area and other cortical areas in organization of voluntary movements in man. Journal of Neurophysiology 43, 118–136. 10.1152/jn.1980.43.1.118

Rolls, E.T., Huang, C.-C., Lin, C.-P., Feng, J., Joliot, M., 2020. Automated anatomical labelling atlas 3. NeuroImage 206, 116189. 10.1016/j.neuroimage.2019.116189

Salardini, A., Narayanan, N.S., Arora, J., Constable, T., Jabbari, B., 2012. Ipsilateral synkinesia involves the supplementary motor area. Neuroscience Letters 523, 135–138. 10.1016/j.neulet.2012.06.060

Schalk, G., McFarland, D.J., Hinterberger, T., Birbaumer, N., Wolpaw, J.R., 2004. BCI2000: a general-purpose brain-computer interface (BCI) system. IEEE Transactions on Biomedical Engineering 51, 1034–1043. 10.1109/TBME.2004.827072

Scherer, R., Müller-Putz, G.R., Pfurtscheller, G., 2007. Self-initiation of EEG-based brain–computer communication using the heart rate response. J. Neural Eng. 4, L23. 10.1088/1741-2560/4/4/L01

Sebastiani, L., Di Gruttola, F., Incognito, O., Menardo, E., Santarcangelo, E.L., 2019. The higher the basal vagal tone the better the motor imagery ability. Arch Ital Biol 157, 3–14. 10.12871/00039829201911

Serino, A., Bockbrader, M., Bertoni, T., Colachis IV, S., Solcà, M., Dunlap, C., Eipel, K., Ganzer, P., Annetta, N., Sharma, G., Orepic, P., Friedenberg, D., Sederberg, P., Faivre, N., Rezai, A., Blanke, O., 2022. Sense of agency for intracortical brain–machine interfaces. Nat Hum Behav 6, 565–578. 10.1038/s41562-021-01233-2

Shibasaki, H., Sadato, N., Lyshkow, H., Yonekura, Y., Honda, M., Nagamine, T., Suwazono, S., Magata, Y., Ikeda, A., Miyazaki, M., Fukuyama, H., Asato, R., Konishi, J., 1993. Both primary motor cortex and supplementary motor area play an important role in complex finger movement. Brain 116, 1387–1398. 10.1093/brain/116.6.1387

Silvani, A., Calandra-Buonaura, G., Dampney, R.A.L., Cortelli, P., 2016. Brain-heart interactions: physiology and clinical implications. Philos Trans A Math Phys Eng Sci 374. 10.1098/rsta.2015.0181

Sitaram, R., Ros, T., Stoeckel, L., Haller, S., Scharnowski, F., Lewis-Peacock, J., Weiskopf, N., Blefari, M.L., Rana, M., Oblak, E., Birbaumer, N., Sulzer, J., 2017. Closed-loop brain training: the science of neurofeedback. Nat Rev Neurosci 18, 86–100. 10.1038/nrn.2016.164

Solomon, J.P., Kraeutner, S.N., O’Neil, K., Boe, S.G., 2021. Examining the role of the supplementary motor area in motor imagery-based skill acquisition. Exp Brain Res 239, 3649–3659. 10.1007/s00221-021-06232-3

Stoodley, C.J., Schmahmann, J.D., 2009. Functional topography in the human cerebellum: A meta-analysis of neuroimaging studies. NeuroImage 44, 489–501. 10.1016/j.neuroimage.2008.08.039

Stoodley, C.J., Valera, E.M., Schmahmann, J.D., 2012. Functional topography of the cerebellum for motor and cognitive tasks: An fMRI study. NeuroImage 59, 1560–1570. 10.1016/j.neuroimage.2011.08.065

Tadel, F., Baillet, S., Mosher, J.C., Pantazis, D., Leahy, R.M., 2011. Brainstorm: A User-Friendly Application for MEG/EEG Analysis. Computational Intelligence and Neuroscience 2011, 879716. 10.1155/2011/879716

Thobois, S., Dominey, P.F., Decety, J., Pollak, P., Gregoire, M.C., Bars, D.L., Broussolle, E., 2000. Motor imagery in normal subjects and in asymmetrical Parkinson’s disease. Neurology 55, 996–1002. 10.1212/WNL.55.7.996

Valenza, G., Di Ciò, F., Toschi, N., Barbieri, R., 2024. Sympathetic and Parasympathetic Central Autonomic Networks. Imaging Neuroscience. 10.1162/imag_a_00094

Valenza, G., Faita, F., Citi, L., Saul, J., Bruno, R., Barbieri, R., 2020. Validation of Sympathetic Activity Index From Heart Rate Variability series: A Preliminary Muscle Sympathetic Nerve Activity Study, in: 2020 Computing in Cardiology. Presented at the 2020 Computing in Cardiology, pp. 1–4. 10.22489/CinC.2020.365

Valenza, G., Sclocco, R., Duggento, A., Passamonti, L., Napadow, V., Barbieri, R., Toschi, N., 2019. The central autonomic network at rest: Uncovering functional MRI correlates of time-varying autonomic outflow. NeuroImage 197, 383–390. 10.1016/j.neuroimage.2019.04.075

Venot, T., Desbois, A., Corsi, M.C., Hugueville, L., Saint-Bauzel, L., De Vico Fallani, F., 2024. Intentional binding for noninvasive BCI control. J. Neural Eng. 10.1088/1741-2552/ad628c

Vidaurre, C., Blankertz, B., 2010. Towards a Cure for BCI Illiteracy. Brain Topogr 23, 194– 198. 10.1007/s10548-009-0121-6

Virameteekul, S., Bhidayasiri, R., 2022. We Move or Are We Moved? Unpicking the Origins of Voluntary Movements to Better Understand Semivoluntary Movements. Front. Neurol. 13. 10.3389/fneur.2022.834217

Wallin, B.G., Fagius, J., 1988. Peripheral Sympathetic Neural Activity in Conscious Humans. Annual Review of Physiology 50, 565–576. 10.1146/annurev.ph.50.030188.003025

Welniarz, Q., Worbe, Y., Gallea, C., 2021. The Forward Model: A Unifying Theory for the Role of the Cerebellum in Motor Control and Sense of Agency. Front. Syst. Neurosci. 15. 10.3389/fnsys.2021.644059

Wolpaw, J.R., McFarland, D.J., Neat, G.W., Forneris, C.A., 1991. An EEG-based brain-computer interface for cursor control. Electroencephalography and Clinical Neurophysiology 78, 252–259. 10.1016/0013-4694(91)90040-B

Wolpaw, J.R., McFarland, D.J., Vaughan, T.M., Schalk, G., 2003. The Wadsworth Center brain-computer interface (BCI) research and development program. IEEE Transactions on Neural Systems and Rehabilitation Engineering 11, 1–4. 10.1109/TNSRE.2003.814442

Wolpert, D.M., Miall, R.C., Kawato, M., 1998. Internal models in the cerebellum. Trends Cogn Sci 2, 338–347. 10.1016/s1364-6613(98)01221-2

Yang, G., Ye, Q., Xia, J., 2022. Unbox the black-box for the medical explainable AI via multi-modal and multi-centre data fusion: A mini-review, two showcases and beyond. Information Fusion 77, 29–52. 10.1016/j.inffus.2021.07.016

Zimmermann-Schlatter, A., Schuster, C., Puhan, M.A., Siekierka, E., Steurer, J., 2008. Efficacy of motor imagery in post-stroke rehabilitation: a systematic review. Journal of NeuroEngineering and Rehabilitation 5, 8. 10.1186/1743-0003-5-8

